# Impact of Sperm Fractionation on Chromosome Positioning, Chromatin Integrity, DNA Methylation and Hydroxymethylation Level

**DOI:** 10.1101/2025.06.07.658443

**Authors:** Zuzanna Graczyk, Jagoda Kostyk, Julia Pospieszna, Zuzanna Myslicka, Marzena Kamieniczna, Monika Fraczek, Marta Olszewska, Maciej Kurpisz

**Affiliations:** Institute of Human Genetics, Polish Academy of Sciences, Strzeszynska 32, 60-479 Poznan, Poland; Poznan University of Medical Sciences, Department of Toxicology, Collegium Pharmaceuticum, Rokietnicka 3, 60-806 Poznan, Poland

**Keywords:** sperm chromosomes, nuclear order, sperm methylation, sperm hydroxymethylation, sperm fractions, sperm DNA fragmentation, sperm chromatin, male infertility, swim up, density gradient

## Abstract

**Background:** Sperm chromosomes are non-randomly organized in the cell nucleus, which plays an important role in the regulation of early embryo development. It is determined by the specific localisation of sperm chromosomal regions carrying genes with expression crucial at the first contact with ooplasm during fertilization. Thus, the aim of this study was to determine whether the application of selective methods providing high-quality spermatozoa with good motility and/or morphology, may increase the frequency of gametes with the specific positioning of chromosomes. For the first time we have used the sequential staining algorithm for consecutive analyses of the same individual sperm cells with a fixed position, what enables to achieve a full and detailed documentation at the single cell level.

**Methods:** Semen samples from 5 normozoospermic males were collected and processed for fractionation *via* swim up (to select viable and motile spermatozoa) or Percoll density gradient (90/47%; for good sperm motility and morphology). Sperm chromatin protamination was assessed by aniline blue (AB) staining, while DNA fragmentation by acridine orange (AO) (ssDNA fragmentation) or TUNEL assay (ssDNA and dsDNA fragmentation). Then, sequential staining and analyses of the same individual sperm cell with a fixed position on a slide were performed, in the following order: (i) fluorescence *in situ* hybridization (FISH) for determination of positioning of chromosomal centromeres: 4, 7, 8, 9, 18, X and Y, with so-called: linear and radial estimations applied, followed by distance measurements between selected pairs of chromosomes; and (ii) immunofluorescent (IF) measurement of global sperm DNA methylation (5mC) and hydroxymethylation (5hmC) levels, what added additional data about epigenetic layer of the sperm chromosomes’ positioning.

**Results:** Our study demonstrated that high-quality sperm selection methods significantly: (i) increased frequency of spermatozoa with good chromatin protamination (+∼25%) and 5mC and 5hmC DNA levels (+∼9.5%), and (ii) reduced rate of spermatozoa with ssDNA fragmentation (-∼65%). Motile and morphologically normal spermatozoa showed distinct chromosome repositioning with sex chromosomes shifted to the nuclear periphery, a key chromosomal region of the initial interaction with the ooplasm during fertilization process. Evaluated autosomes revealed various patterns of repositioning.

Conclusions

Our findings underline the validity of methods used for selection of high-quality spermatozoa in assisted reproductive technologies (ART), also in the context of the sperm chromosomal topology and chromatin integrity, crucial at the first steps during fertilization.

## Introduction

Infertility is a significant global health issue, affecting approximately 10-18% of couples in the reproductive age, with male background contributing to 40-60% of all cases (1). Male infertility can result from various environmental factors, including: tobacco smoking, drug use, anabolic steroids, alcohol consumption, followed by exposure to: heavy metals, radiation, organic chemicals, or high temperatures (2). Genomic and cytogenomic aberrations account for approximately 10-15% of male infertility cases (3–5). What is important, approximately 50% of genetic causes is strictly linked to spermatogenesis disruptions, underlining the high genetic vulnerability of the process of male gametes generation – spermatozoa with a haploid genome. Their task is to deliver to the oocyte the paternal genetic material important for the development of the embryo (6,7).

The chromatin organization of spermatozoa differs from chromatin packaging in somatic cells. Mainly, during spermatogenesis, the sperm chromatin becomes drastically compacted and also transcriptionally silenced at an extremely high level (8–10). This chromatin rearrangement occurs because of an exchange of at least 85% of the somatic type of histones: first, by transition proteins (TPs), and then by the protamines, leading to a formation of DNA-protamine complexes (toroids) with strongly condensed chromatin – about 6-fold more compact than that in somatic cells (8,11,12). This unique chromatin organization observed in the sperm nucleus likely serves several critical functions, including: (i) the efficient compaction of paternal DNA, to protect the genome from potential damage while passaging through the female reproductive tract (13); (ii) effective mass silencing of sperm gene expression, but with preserved capacity for rapid reactivation of genes following fertilization (14); (iii) efficient repair of DNA damage and remodelling of the paternal chromatin, critical for early embryonic divisions through the sequential order of the paternal genome delivering to the maternal oocyte (8,15). It has been also clearly shown, that abnormalities in histone transition and protamine amount in humans may be associated with defective spermatogenesis leading to decreased sperm quality, possible male infertility, and failed attempts of *in vitro* fertilization by assisted reproductive technology (ART) (9,10,16–18).

It has been documented that the location of the chromosomes in cell nucleus is well-defined and non-random within the so-called: ‘chromosome territories’ (CTs). CTs together with topologically associating domains (TADs) and interchromatin compartments (ICs) interact with the elements of the nuclear matrix creating the intranuclear architecture (19–22). Therefore, the topology of chromosomes may have specific characteristics by determining location of their centromeres and/or the CTs, as well as the p and q arms of chromosomes (22). As for somatic cells there is a variety of novel techniques with ultra-high resolution (i.e., GAM, Hi-C, SPRITE, ChIA-Drop) for exploring TADs and all possible linkages between particular regions of the chromosomes (15,22). In case of spermatozoa the visualization of chromosomes still relies on fluorescent *in situ* hybridization (FISH) with probes specific for particular parts of chromosome (arms, centromeres, bands, (sub)telomere regions, etc.) as sperm cells remain virtually transcriptionally inactive, limiting the applicability of techniques that depend on RNA or active transcriptional processes (12,15,23,24). Determination of the organization of chromosomes in the human sperm cell nucleus is based mostly on the linear and radial localisation of centromeres (8,11,12,23,25,26) that allow to estimate their positioning in a preferential part of the nucleus, in relation to the sperm acrosome, its centre or tail region, as well as the localization’s depth in a 3D manner.

It was also suggested that the sperm cell nucleus architecture plays an important role in the regulation of early embryo development via the first contact with ooplasm of the chromosomal regions carrying specific genes whose expression is crucial at the first stages of embryo development (6,15,25). For instance, in human sperm X chromosome is typically positioned within the subacrosomal region of the sperm nucleus (12,27–29). This localization may be important for rapid activation of genes immediately after the fusion of gametes. A subset of genes located on sex chromosomes (e.g. *RPS4Y1* (Gene ID: 6192) and *RPS4X* (Gene ID: 6191)) are transcription or epigenetic factors with downstream autosomal targets, and are expressed soon after fertilization (30–34). Given their regulatory roles, it is likely that sex chromosomes are transcribed first, as they influence the expression of autosomal genes (e.g. *KDM6A* (Gene ID: 7403)*, SRY* (Gene ID: 6736)) essential for initiating proper regulatory networks and ensuring correct embryonic development (30,31,34). The development of sex-specific regulatory networks enriched in X- and Y-linked genes plays a critical role in cellular differentiation, also during early mammalian development (35–37). Moreover, female embryos undergo X chromosome inactivation (XCI) during implantation, to balance the X-linked gene dosage between XY and XX embryos (38–44). Impaired XCI is one of the major epigenetic barriers, that can hinder correct development of female embryos, often resulting in early miscarriage and embryonic lethality (45–47). In humans, after the completion of embryonic genome activation at E4, female cells possess two active X chromosomes (48). Both of X chromosomes in females are widely activated immediately after embryo genome activation from the 4- to the 8-cell stage. What is important, distinct activation patterns of sex chromosomes are observed during early embryogenesis, with only a few genes on the Y chromosome being activated initially and with activity of a broad region on the X chromosome (34). For example, *RPS4Y1* (Gene ID: 6192), a gene located on Y chromosome, is highly expressed during embryonic genome activation and shows a sex-specific pattern. Its paralog, *RPS4X* (Gene ID: 6191), is present on a long arm of the X chromosome (Xq) (49). *RPS4X* is also known to escape from X inactivation (49). It has been assumed that normal human development requires two *RPS4* genes per cell: two copies of *RPS4X* in female cells, and one copy of *RPS4X* and one copy of *RPS4Y* in male cells (34). The high transcription of *RPS4Y1* helps balance the gene dosage between sexes during early development, compensating for the two-fold dosage of *RPS4X* in females after embryonic genome activation, as both X chromosomes remain active (50).

Another example supporting the importance of sperm cell nucleus architecture, as well as the observation that X chromosomes are typically positioned within the subacrosomal region of the sperm nucleus, which may facilitate the rapid activation of genes immediately after gamete fusion, is the *SMC1A* gene (Gene ID: 8243) located on the X chromosome, in an area that escapes XCI (51). *SMC1A* encodes a structural component of the cohesin complex, which ensures proper cohesion of sister chromatids in mitosis and meiosis (52). This is crucial for the correct segregation of chromosomes during cell division. The encoded protein is also thought to be an important part of functional kinetochores (51). In addition, this protein has a potential role in DNA repair and genome stability maintenance (53–56). Furthermore, together with *CTCF* transcription factor (Gene ID: 10664), *SMC1A* takes part in organizing the three-dimensional structure of the genome also in pre-implantation embryo development (57–60).

These examples seem to highlight the significance of X- or Y-located genes delivered by sperm cells, underlining the importance of the sex chromosomes’ spatial organisation within the sperm nucleus. Especially in the fact, that activation of sex chromosome-linked regulatory networks are known to be critical for early embryonic development and cellular differentiation (34,61). Adapting of ultra-high-resolution techniques like GAM or Hi-C for use in evaluation of sperm cells could significantly improve our understanding of the unique organization of the sperm cell nucleus. However as mentioned above, the almost transcriptionally inactive nature of sperm cells limits the applicability of methods relying on RNA or active transcriptional processes, which explains the current reliance on FISH for chromosomal visualization (62,63). However, a promising direction for future studies in the field is implementation of such cutting-edge techniques in sperm research would help uncover the specific interactions between chromosomal regions and would shed light on how nuclear architecture contributes to the regulation of early embryonic development.

Another issue crucial both for sperm characteristics, as well as for fertilization or early embryo development events are epigenetic modifications of sperm DNA or histones (58,64–68). It was found that sperm DNA methylation (5mC) and DNA hydroxymethylation (5hmC) can affect the results of ART and is linked with embryonic development (69,70). Sperm cells which in principle are transcriptionally inactive, exhibit distinct methylation patterns characterized by global hypermethylation of sperm DNA in contrast to hypomethylated somatic cells. This hypermethylation is predominantly observed in intergenic regions and repetitive elements, contributing to support genomic stability (66). The appropriate parent-of-origin expression in the developing embryo strictly relies on the hypermethylation of imprinted genes (13,67,71,72). On the other hand, promoters of developmental genes and imprinting control regions are specifically hypomethylated, which is essential for proper gene expression during early embryonic development. This hypomethylation facilitates the instant activation of developmental genes, leading to epigenetic reprogramming and developmental assignment determination (13,58,65,67,68,73,74). Understanding these sperm-specific epigenetic patterns and their regulatory consequences is crucial for elucidating the mechanisms of transgenerational epigenetic inheritance and their impact on offspring health. Recent studies have shown that environmentally induced parental epigenetic alterations can be transmitted into the next generations and influence the phenotype of the offspring and their potential disease development (75–79). When focusing on DNA methylation, it is well-established that it is crucial for processes such as genetic imprinting, gene silencing, chromosome X inactivation and protein conformational changes (67,71,73,80–82). Parental genomes have distinct genetic roles after fertilization, driven by gametic imprinting and unique methylation patterns established during gametogenesis (13,67,71,72,83). The paternal genome primarily supports early placental development, while the maternal genome governs embryonic development (13,67,72,83). Developmental disturbances in embryos may arise from improper activation of key genes, often linked to disrupted methylation/demethylation cycles in gametogenic cells (13,67,68,74,84–86). Hydroxymethylation (5hmC) occurs at lower levels (0.1–0.8%) compared to methylation, with higher values in tissues with active transcription, such as neurons (82,87–89). This epimark, derived from enzymatic oxidation of 5mC, is found in enhancers, gene promoters, and regulatory elements (82,90). In transcriptionally inactive spermatozoa, 5hmC levels are about 4-fold lower than in somatic cells (91). Together with Tet enzymes (Ten-Eleven Translocation Proteins; Tet1, Tet2, Tet3), 5hmC may regulate gene expression by modulating methylation, highlighting the critical roles of both 5mC and 5hmC in genome function (87,89,91–94).

Sperm fractionation is routinely utilized in the infertility treatment clinics for assisted reproductive technologies (ART) such as *in vitro* fertilization (IVF) or intracytoplasmic sperm injection (ICSI). Some of the well-known methods of sperm separations are swim-up fractionation or density-gradient centrifugation (69). Swim-up method allows to separate motile fraction of spermatozoa from seminal plasma, while density-gradient centrifugation, (i.e., Percoll gradient), allows to separate morphologically normal and motile human spermatozoa, free of debris, dead cells and non-germ cells (1). Sperm cells obtained by these two methods result in similar cumulative life birth rate (95). Evidence showed that the selection of spermatozoon before injection into the oocyte has a significant impact on ICSI outcomes (96). Choosing proper spermatozoa remains a pivotal stage, influencing fertilization processes and early embryo development. Nevertheless, the main challenge arises from the varying quality of spermatozoa within an ejaculate (97). The *zona pellucida* of the oocyte acts as a selective barrier during fertilization and prevents penetration of multiple spermatozoa. During intracytoplasmic sperm injection (ICSI or IMSI – Intracytoplasmic Morphologically Selected Sperm Injection), the best spermatozoa for oocyte insemination are selected according to their correct shape and motility (98). However, such visual inspection for normal spermatozoa does not eliminate the genetically abnormal spermatozoa – the visually normal sperm cells can be (and often are) carriers of genetic or epigenetic mutations that may disturb fertilization events, embryo development or be devastating for the offspring. Therefore, it is important to deepen the knowledge of the mechanisms influencing sperm quality to improve the selection process for broadly used *in vitro* fertilization.

In this context, the aim of this study was to investigate for the first time whether chromosomal localization differs good-quality fractions of human spermatozoa separated via two different protocols: swim-up or Percoll density gradient, followed by sperm DNA methylation and hydroxymethylation levels screening. This aim was achieved by sequential staining algorithm of the same individual sperm cell with a fixed position on a slide, in an given order: (i) determination of chromosomal positioning of centromeres: 4, 7, 8, 9, 18, X and Y, then (ii) measurement of sperm DNA 5mC and 5hmC levels. Results were additionally supported by sperm chromatin integrity assays. The knowledge obtained by implementation of our sequential staining clearly reveals the validity of application of sperm separation methods regarding the chromosomal context of fertilization.

## Materials and methods

### Participants

Biological material consisted of sperm cells from n=5 healthy male normozoopermic volunteers (according to WHO criteria (1)) aged between 25 to 30 years, with 46,XY karyotype, and without history of reproductive failure. All men were notified about the purpose of the study and provided written informed consent, according to the approved protocols and guidelines of the Local Bioethical Committee at Poznan University od Medical Sciences (approval no. 669/22).

Schematic workflow of experimental approaches in this study, including sperm chromatin quality check and sequential staining algorithm is presented in Figure 1.

**Figure 1.**
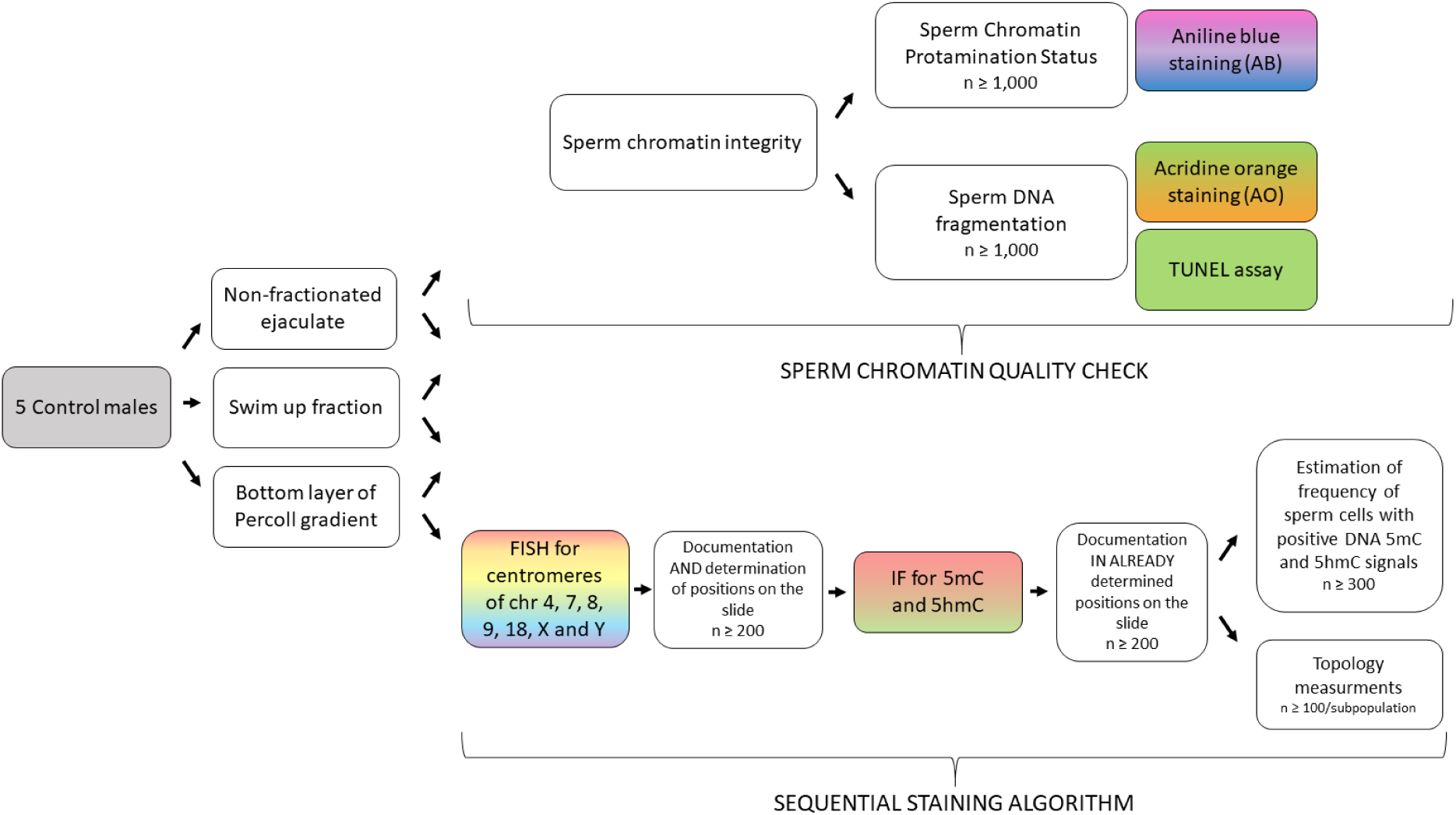
Workflow chart of the experimental approach. Step by step experimental algorithm included separation of good quality fractions, and then two groups of experiments. First, sperm chromatin quality check (upper panel) has been done to characterize the group of evaluated cases. Second, sequential staining algorithm (lower panel) involved: (i) the determination of chromosomal positioning (centromeres of chromosomes: 4, 7, 8, 9, 18, X, and Y) via fluorescence *in situ* hybridization (FISH), and (ii) the subsequent analysis of global DNA methylation (5mC) and hydroxymethylation (5hmC) levels in the same sperm cells. n = number of sperm cells measured for each of the evaluated chromosome.

### Preparation of semen samples

Semen samples were collected *via* masturbation, after 3-4 days of sexual abstinence. After liquefaction (30 min.) samples were analysed manually according to the WHO 2021 (1) criteria for semen evaluation (volume, concentration, motility, morphology, and viability) (Table 1). Then, samples were subjected to fractionation via swim up (to select motile spermatozoa) or Percoll density gradient (to select spermatozoa fraction enriched in good motility and morphology cells), fixed and stored in -20°C until further use. Also, non-fractionated ejaculate samples, washed out from seminal plasma, have been fixed as further described.

**Table 1.** Characteristics of semen parameters of n=5 cases evaluated in the study.

| Case | Age | Sperm concentration (10<sup>6</sup>/ml) | Ejaculate volume (ml) | Total sperm count (x10<sup>6</sup>) | Sperm morphology (normal forms; %) | Total motility (%) | Progressive motility (%) |
| --- | --- | --- | --- | --- | --- | --- | --- |
| **K1** | 30 | 101.23 | 3.0 | 303.69 | 21 | 76 | 59 |
| **K2** | 26 | 48.93 | 2.5 | 122.3 | 4 | 74 | 66 |
| **K3** | 30 | 82.0 | 4.5 | 369.0 | 7 | 63 | 55 |
| **K4** | 25 | 80.25 | 3.8 | 305.0 | 4 | 57 | 48 |
| **K5** | 30 | 122.0 | 2.0 | 244.0 | 6 | 67 | 62 |
| Mean | 28.2 | 86.88 | 3.16 | 268.8 | 9.5 | 67.4 | 58 |
| SD | 2.49 | 27.14 | 1.00 | 93.07 | 7.77 | 7.83 | 6.89 |
| Lower reference limits for normozoospermia according to WHO guidelines (1) | - | >15 | ≥1.5 | >39 | >4 | >40 | >32 |

### Swim up selection

A 1 ml of liquefied semen sample was placed in a round-bottom tube (15 ml; Falcon, Corning, Tamaulipas, Mexico) and centrifuged (8 min., 1800 rpm) with warm (37°C) F10 medium (cat. no. 134464606, Biomed-Lublin, Poland) to remove seminal plasma. The supernatant was discarded, and fresh, warm medium (37°C, 2 ml) gently overlayered the pellet. After incubation (37°C, 45 min.), two fractions were separated: upper layer with mostly motile sperm cells, and the bottom one with non-motile gametes. For aimed examinations top good-quality fractions were selected, named as ‘Swim up fraction (SU)’ for study purposes. Then, samples were subjected to three rounds of fixation in a fresh fixative solution (methanol:acetic acid, 3:1 v/v, -20°C, POCh, Gliwice, Poland), and stored in -20°C until further use.

### Percoll gradient centrifugation

The gradient of 47%/90% of Percoll (Sigma-Aldrich, St. Louis, MO, USA, P1644) was used for separation of spermatozoa with good motility and morphology (99). At the top of the gradient a 1 ml of liquefied semen sample was carefully added and centrifuged in 1495 rpm for 30 min., RT. For the purpose of the study, we selected enriched motile and good morphology spermatozoa contained in bottom fraction of Percoll called for the study purpose as: ‘bottom layer of Percoll gradient (BPG)’. Each sample was washed 3 times in 1x PBS (Gibco, Paisley, UK, pH 7.4) to deplete any traces of Percoll solution, followed by three rounds of fixation (methanol:acetic acid, 3:1 v/v, -20°C). Fixed samples were stored in -20°C until further use.

### Sperm Chromatin Integrity

Sperm chromatin integrity status was evaluated using three tests according to previously published protocols (23,26,100–104). To determine sperm chromatin protamination level the aniline blue (AB) staining was used. To evaluate sperm DNA fragmentation, the acridine orange (AO) staining was performed for sperm single-stranded DNA (ssDNA) fragmentation level estimation, followed by TUNEL assay for determination of sperm double-stranded (dsDNA) and single-stranded DNA (ssDNA) fragmentation level.

### Sperm Chromatin Protamination Status

Sperm chromatin protamination status was evaluated by application of aniline blue (AB) staining. Briefly, fixed sperm cells were spread onto microscopic slides, washed in 2x SSC (saline sodium citrate; pH 7.2, W302600, Sigma-Aldrich) for 3 min. and air-dried. Next, 100 µl of 1% eosin-Y solution (Merck, Darmstadt, Germany) was applied on the slides for 3 min. incubation, followed by water rinsing. Then, slides were stained in acidic 5% solution of aniline blue (cat. no. 95290, Water blue, Fluka, Darmstadt, Germany) for 5 min., rinsed off with water and air-dried. The slides were analysed using a light microscope (Leica DM5500, 100x oil immersion objective; LAS X software; Germany). In each sample at least n≥1,000 spermatozoa were evaluated. Aniline blue is a reagent that binds to lysine residues in histones, resulting in dark blue staining. Three populations of spermatozoa can be distinguished: pink – sperm cells with proper protamines to histones ratio, purple – sperm cells with disturbed protamines to histones ratio (called ‘semiprotaminated’ for the purpose of this study) and navy blue – deprotaminated spermatozoa with predominant number of remaining histones.

### Sperm DNA fragmentation

Sperm DNA fragmentation status was evaluated using two methods: (i) an acridine orange (AO) test to quantify single-stranded (ss) DNA breaks and (ii) TUNEL assay to estimate the level both of double-stranded (ds) and single-stranded (ss) breaks of DNA. For both tests, sperm cells were analysed and documented using fluorescence microscope (Leica DM5500, Germany; 63x oil immersion objective and filters: DAPI, SpG, SpO, triple BGR) and LAS X software (Leica).

An AO test was carried on slides with fixed sperm cells, as described previously (100). Briefly, dry slides were stained with freshly prepared acridine orange solution: 10 ml of 1% AO (cat. no. 235474, Sigma-Aldrich) in distilled water added to a mixture of 40 ml 0.1 M citric acid (cat. no. 251275, Sigma–Aldrich) and 2.5 ml 0.3 M Na2HPO4x7H2O (cat. no. S9390, Sigma–Aldrich; pH 2.5). After incubation in the darkness (5 min.), the slides were rinsed off with distilled water and air-dried. The coverslips were applied on the slides and in each sample at least n≥1,000 spermatozoa were evaluated. Acridine orange test detects and quantifies ss breaks of DNA by utilizing a reagent that binds to DNA and then forms two types of complexes: A and B. The type A complex, as a result of intercalation of monomers between bases in double-stranded DNA, exhibits a maximum absorption at a wavelength of 502 nm (green fluorescence). The type B complex is formed when acridine orange molecules aggregate on a single strand of denatured DNA. The maximum absorption for this complex is 475 nm (red or yellow fluorescence). Thus, two populations of spermatozoa can be distinguished: green – without ssDNA fragmentation and yellow or red – with ssDNA fragmentation.

The TUNEL assay was carried out according to the manufacturer’s protocol, using the FlowTACS Apoptosis Detection Kit (cat. no. 4817-60-K, R&D Systems, Minneapolis, MN, USA) to identify sperm cells with fragmented DNA (presence of nicks) by creation of a complex between biotinylated DNA fragments and streptavidin-conjugated fluorescein (FITC) in the presence of terminal deoxynucleotidyl transferase (TdT) (26,100–102,104). Briefly, after permeabilization in 0.1% Triton/sodium citrate solution (15 min. RT), slides were washed with 1x PBS (Gibco, Paisley, UK, pH 7.4), followed by incubation with TdT and a labelling buffer (1 h at 37°C in the dark). Next, the slides were washed with 1x PBS twice and air-dried. The slides were counterstained with 15 µl of DAPI. TUNEL-positive cells (with fragmented DNA) were fluorescently labelled (green colour), then visualised and counted under a fluorescence microscope. In each sample at least n≥1,000 spermatozoa were evaluated.

After quality estimation of semen parameters and sperm chromatin described above, the sequential staining algorithm has been implemented, as schematically presented in Figure 1.

### Fluorescence in situ hybridization (FISH)

Slides with fixed sperm cells were incubated in a decondensation solution (15 mmol dithiothreitol/ddH_2_O (DTT; cat. no. 111474, Merck KGaA, Darmstadt, Germany), at 43°C for 7-8 min., a chromatin loosening step is required for hybridization of FISH probes to DNA. Then, the slides were rinsed in 2x SSC (pH 7.2) and air-dried. The volume of the sperm nucleus increased to 1.4-1.5-fold, maintaining shape and geometrical features of the spermatozoon. The FISH technique was prepared according to the manufacturer’s protocol (Cytocell, Cambridge, UK) with minor modifications (23,103,105). Briefly, slides with decondensed sperm cells were washed in 2x SSC (3 min.), dehydrated with ethanol series (70-85-99%) and allowed to dry. To investigate the localization of centromeres of chromosomes: 4, 7, 8 and 9, 2.5 µl of each α-satellite or satellite III probe (Cytocell, UK) was applied (Additional file 1). The final mix volume was adjusted to 10.0 µl using hybridization buffer. To investigate the localization of centromeres of chromosomes 18, X and Y, 7.0 µl of 18 centromere probe, and 2.5 µl of X and Y probes each (Cytocell, UK) were used, with the hybridization buffer adjustment to 20 µl (Additional file 1). The slides and the mix of probes were pre-warmed at 37°C for 5 min. After that, the probe mixture was placed on a slide and covered with coverslip (24 x 24 mm), sealed with a rubber glue, and denatured in 75°C for 2 min. The slides were placed in a humid, lightproof container and incubated overnight at 37°C for hybridization. Next, the coverslip was removed and the slide was immersed in: 0.4x SSC (pH 7.0) at 72°C for 2 min., and 2x SSC, 0.05% Tween-20 (pH 7.0) at RT for 30 s, to get rid of any remaining unbound probes. Then, DAPI/antifade was applied on each slide and covered with coverslip (24 x 60 mm). The colour was allowed to develop in the darkness for 10 min.

FISH results were scored using fluorescence microscope (Leica DM5500, Germany; 63x oil immersion objective, DAPI, SpG, SpO, SpA, triple BGR filters and motorized stage). The acquisition of images was performed with LAS X software (Leica, Germany) with Navigator functions that allowed to document the position of the particular sperm cell on the slide and use the same localization for the next round of staining in the sequential staining algorithm.

### Localization of the centromeres

The linear and radial positioning of chromosomes 4, 7, 8, 9, 18, X and Y was estimated as it was developed by Zelenskaya and Zalensky (12). Due to the geometrical features of the spermatozoa on the microscopic slide, developed measurements patterns allow the results to be normalized and located within the nuclear space, ensuring consistent and reliable comparisons across samples. Those patterns have been explained and successfully used in our previous topology studies (23,26,100,103–110).

### Linear positioning of the centromeres of chromosomes

The linear positioning demonstrates the frequency of FISH signals in three equal territories of the sperm nucleus determined along its longitudinal axis: ‘a’ – near the acrosome, ‘m’ – middle part, and ‘t’ near the tail (Fig. 2a).

**Figure 2.**
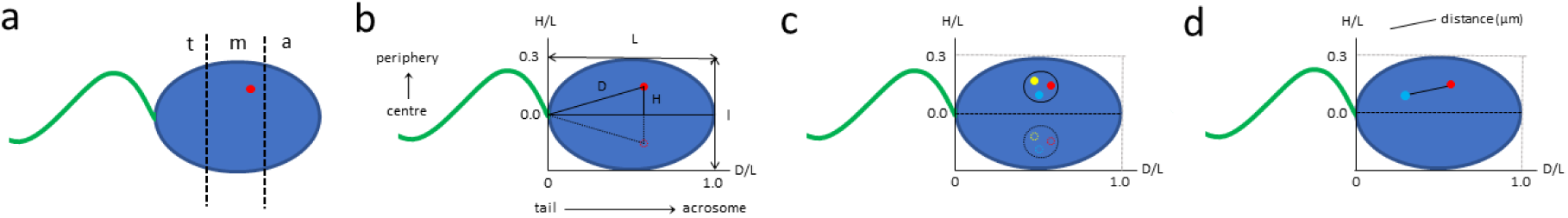
Schematic representation of measurement approaches of centromere localization (red, yellow and light blue points) within sperm nucleus (blue background). **a** Linear positioning: frequency of FISH signals across three equally divided regions of the nucleus, segmented along its longitudinal ’L’ axis (‘t’ – near the tail, ‘m’ – middle, ‘a’ – near the acrosome). **b** Radial positioning: shown as central (deep inside the nucleus – values close to 0.0) or peripheral (near the nuclear membrane – values close to 0.3) localization of the signal, based on normalized D/L (OX axis) and H/L (OY axis) values; D and H – the distances from the FISH signal (red point) to the tail attachment point and the longitudinal axis, respectively; L and l – the longitudinal and short axes, respectively. Dotted lines show mirror images of centromere positioning (the sperm cell has the property that it can only take two positions on the microscope slide, which are mirror images of each other). **c** Schematically marked chromocenter region (clusters of centromeres). **d** The distance (μm) between pair of centromeres within the sperm nucleus. The idea of radial positioning was for the first time introduced by Zalenskaya and Zalensky (12).

### Radial positioning of the centromeres of chromosomes

To determine radial position of chromosomes 4, 7, 8, 9, 18, X and Y in spermatozoa the following measurements of the sperm nucleus dimensions were performed: L – length of the longitudinal axis (from the tail attachment point to the acrosome), l – length of the short axis (in the widest part of the nucleus), D – distance from FISH signal to the tail attachment point, H – distance from FISH signal to the longitudinal axis, and ratio: L/l – the ellipsoidal shape indicating the decondensation ratio of the nucleus (not exceeding 1.4-1.5-fold). The D/L value was used to determine the central localization, towards the ‘tail-acrosome’ direction, with a maximum value of 1.0. The H/L value defined the peripheral localization (‘centre-periphery’ criterion, the proximity to the internal sperm nuclear membrane), with a maximum value of 0.3. The measurement system allows to depict centromeres’ positions in a coordinate system as the mean D/L ± SE for the OX axis and H/L ± SE for the OY axis which allows the results to be normalized and located in the nuclear space (Fig. 2b). By normalizing the obtained data, results can be positioned in a nuclear space that approximates a 3D model, where the relationship between the central and peripheral regions is effectively visualized (12,103). This spatial representation can visualize centromere localization both deeply in the centre of the sperm nucleus or near the nuclear membrane. For each evaluated case, each chromosome, and for each 5mC and 5hmC groups, at least n≥100 FISH signals were measured, which resulted in the analysis of 14,120 sperm cells, in total. The clusters of intranuclear positions of the analysed centromeres were presented as specific regions within the sperm nucleus, referred to as chromocenters (Fig. 2c).

### Distances between the centromeres of chromosomes

For each sample, the distance between the centromeres of two chromosomes was measured for at least n≥100 spermatozoa. The measurement of the distance between centromeres was performed for following pairs of chromosomes: 4 vs. 8, 7 vs. 9, and 18 vs. X or Y (Fig. 2d).

#### 5mC and 5hmC levels of sperm DNA

On slides with previously defined spermatozoa’s position and documented topology of centromeres, immunofluorescence (IF) staining for determination of the global sperm DNA methylation (5mC) and hydroxymethylation (5hmC) levels was performed. This method has been validated previously when correlated to thin-layer chromatography results (TLC) and used with further success (101,102). Specific antibodies conjugated to fluorochromes and diluted in 1%BSA/1xPBST were used: primary antibodies – mouse anti-5mC 1:200 (clone 33D3, cat. no. MABE146, Merck), and rat anti-5hmC 1:1000 (cat. no. ab106918, Abcam); secondary antibodies – goat anti-mouse-FITC 1:400 (cat. no. F2012, Sigma-Aldrich), and goat anti-rat-AF594 1:800 (cat. no. ab150160, Abcam). First, samples with fixed sperm smears after FISH staining and documentation were incubated in 1x PBST (pH 7.4) for 10 min., followed by 4 washes in 1x PBST (pH 7.4, 5 min. each). Then, slides were incubated in 25mM DTT/1M Tris-HCl (pH 9.5, 10 min.) to slightly decondense the chromatin, followed by two washes in 1x PBST (pH 7.4, 5 min. each). To denature the samples, slides were then incubated in: 6 N HCl (15 min.), 1M Tris-HCl (pH 9.5, 15 min.), and 1x PBST (pH 7.4, 5 min.). Next, slides were blocked with 1% BSA/1× PBST for 1h, and incubated overnight in a humidified container at 4°C with a mix of primary antibodies. After incubation and series of rinsing in 1x PBST (threefold, 5 min. each), secondary antibodies conjugated to selected fluorochromes were applied for 1h. Next, samples were washed twice in 1x PBST (pH 7.4, 5 min. each), and DAPI/antifade solution (Vectashield, cat. no. H-1000, Vector Laboratories, Newark, CA, USA) was applied on each slide and covered with coverslip (24 x 60 mm). The colours were allowed to develop in the darkness for 10 min. Results were analysed using fluorescence microscope (Leica DM5500, Germany; 63x oil immersion objective, DAPI, SpG, SpO, triple BGR, and motorized stage). The acquired images were analysed using and LAS X software and Navigator options (Leica, Germany).

As mentioned above, spermatozoa were collected in three groups: non-fractionated ejaculated sample, swim up fraction (SUG) or bottom layer of Percoll gradient (BPG). SUG and BPG were the sperm good-quality fractions. For each fraction and each chromosome, position on slide of at least n=200 spermatozoa were documented. Then, second round of staining for the same spermatozoa with fixed positions (on slides) were performed for estimation of global 5mC and 5hmC levels of sperm DNA. Topology measurement for each case was performed in total of approximately n=4,200 of spermatozoa (7 chromosomes x 3 fractions x 5mC/5hmC hyper/hypo states x at least 100 signals each time) (Fig. 1, Fig. 2). Quantitative immunofluorescence analysis showed that spermatozoa exhibiting high levels of 5mC also tended to display increased 5hmC IF intensity, indicating a correlation between these two DNA modifications within the same cells (111). Therefore, we focused our topology measurements on spermatozoa with either high or low levels of both epimarks.

### Statistical analysis

For verification of the normal distribution of the measurements, the Shapiro-Wilk test was performed. For statistical analysis of sperm chromatin integrity, linear results and estimation of the 5mC and 5hmC levels, Mann Whitney test or unpaired t-test with Welch’s correction were applied with a significance level of α=0.05. For statistical analysis of the radial results and distances between the centromeres of chromosomes, Kruskal-Wallis test with Dunn’s multiple comparisons test was carried out at a significance level of α=0.05. All the tests were performed using GraphPad Prism (v.8.4.3) software. Estimation of the common aggregation of centromeres was performed using Ward cluster analysis and visualized as hierarchical trees with linkage to Euclidean distances, utilizing the tool available on the website https://datatab.net/statistics-calculator/cluster.

## Results

The results were presented in two sections: first, the assessment of sperm quality parameters, including chromatin protamination status, DNA fragmentation, and global DNA methylation and hydroxymethylation levels (5mC and 5hmC); followed by the analysis of chromosomal positioning and epigenetic modifications performed by sequential staining and evaluation of the same individual sperm cells.

### Sperm Chromatin Integrity

The results of sperm chromatin integrity evaluation consisted of sperm chromatin protamination test (AB) and sperm DNA fragmentation assays (AO and TUNEL) were presented in: Table 2, Figure 3a, Figure 4 and Additional file 2

**Figure 3.**
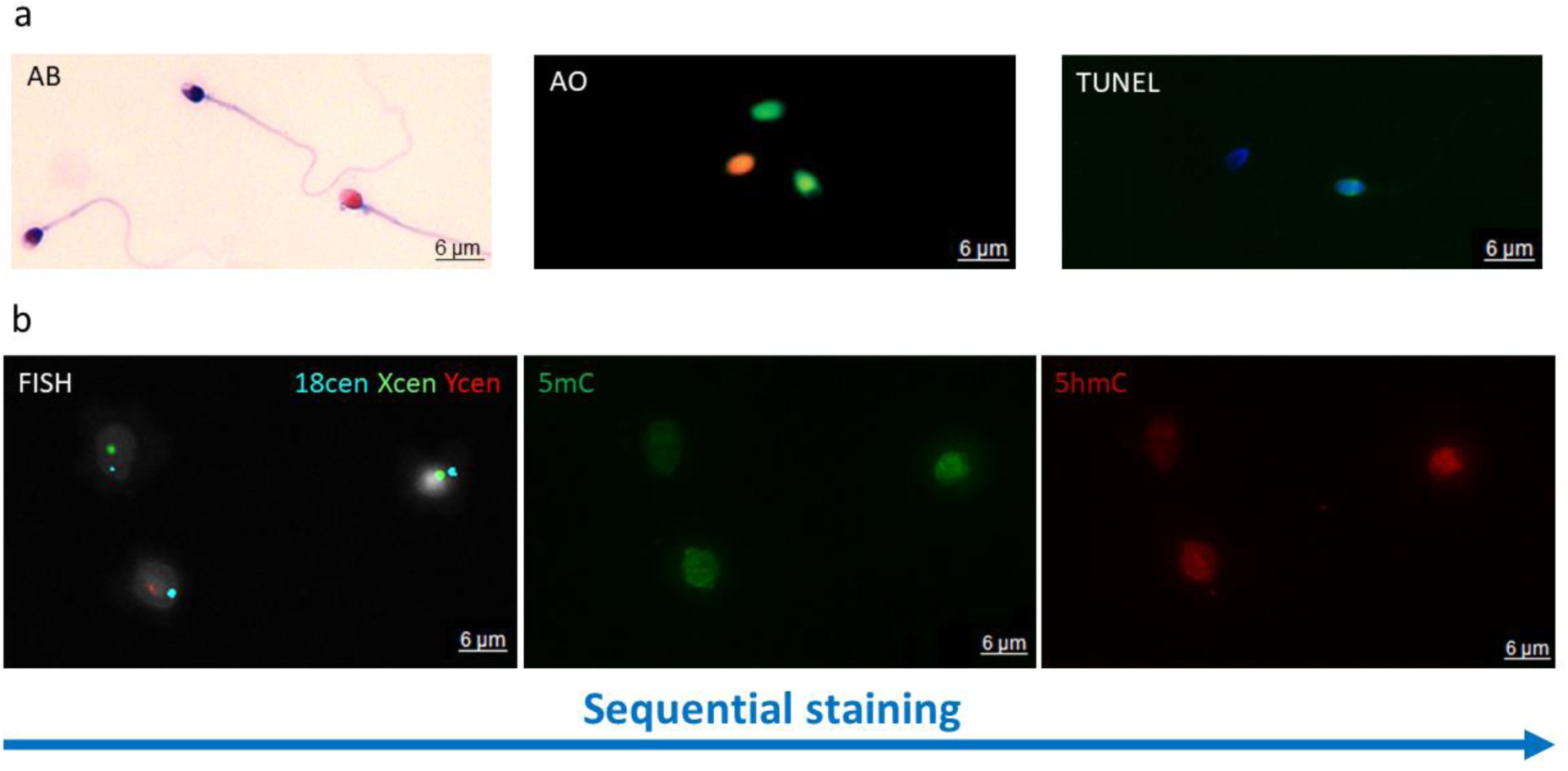
Examples of staining results. **a** Sperm chromatin quality check: (i) Aniline blue (AB) staining: three different sperm subpopulations: pink with proper chromatin protamination level, purple with disturbed protamination, and navy blue with highly deprotaminated chromatin, (ii) Acridine orange (AO) staining: two different sperm subpopulations: green without ssDNA fragmentation and orange with ssDNA fragmentation, (iii) TUNEL assay: two sperm nuclei subpopulations: TUNEL- negative (without dsDNA and ssDNA fragmentation) and TUNEL-positive bright green spermatozoa (with dsDNA and ssDNA fragmentation). **b** Step-by-step sequential staining algorithm of spermatozoa. First – fluorescence *in situ* hybridization (FISH) for centromeres of selected chromosomes (left panel); second – immunofluorescent staining (IF) for evaluation of global levels of sperm DNA methylation (5mC) and hydroxymethylation (5hmC) (middle and right panels). Images were acquired with light (AB) or fluorescent (AO, TUNEL, FISH, IF) microscope (Leica DM5500, oil immerse objective 63x and SpO/FITC/SpAq/DAPI filters).

**Table 2.** Results of sperm chromatin integrity evaluation in non-fractionated sperm population and good-quality fractions (swim up fraction (SU) and Percoll gradient (BPG))

| | | Non-fractionated sperm<br>Mean ± SD | SU<br>Mean ± SD | BPG<br>Mean ± SD |
| --- | --- | --- | --- | --- |
| Sperm chromatin protamination [%; aniline blue assay, AB] | Properly protaminated (pink) | 75.47 ± 9.51 | 93.76* ± 2.75 | 95.04* ± 3.98 |
| | Semiprotaminated (purple) | 13.77 ± 6.95 | 3.22* ± 1.69 | 3.39* ± 3.29 |
| | Deprotaminated (navy blue) | 10.76 ± 9.03 | 3.02 ± 1.22 | 1.57 ± 0.84 |
| | Sum of semi- and deprotaminated | 24.53 ± 9.51 | 6.24* ± 2.75 | 4.96* ± 3.97 |
| Spermatozoa with fragmented DNA [%] | Spermatozoa with fragmented ssDNA [Acridine Orange staining, AO] | 21.03 ± 7.04 | 6.70* ± 7.96 | 6.39** ± 3.40 |
| | Spermatozoa with fragmented ssDNA and dsDNA [TUNEL assay] | 5.54 ± 2.53 | 5.50 ± 3.94 | 2.76 ± 2.18 |

\* values statistically significant ( $p < 0.05$ ) from the non-fractionated sperm values;
\* values highly statistically significant ( $p < 0.01$ ) from the non-fractionated sperm values;

The results of aniline blue (AB) staining showed that the average level of sperm DNA protamination was 75.47 ± 9.51% (range: 65.42-77.5%) in non-fractionated ejaculate, 93.76 ± 2.75% (range: 89.54- 96.94%) in spermatozoa with good motility (swim up fraction, SU) and 95.04 ± 3.98% (range: 87.97- 97.22%) in spermatozoa with good motility and morphology (bottom layer of Percoll gradient, BPG). Thus, the average level of sperm DNA protamination increased in high quality selected spermatozoa: 1.24-fold (+24.24%) for SU sperm fraction (p=0.0105) and 1.3-fold (+25.93%) for BPG sperm fraction (p=0.0159) (Table 2 and Figure 4).

Results of acridine orange (AO) staining showed that the mean frequency of sperm with ssDNA fragmentation was 21.03 ± 7.04% (range: 10.47-29.55%) in non-fractionated spermatozoa, 6.70 ± 3.94% (range: 1.27-20.55%) in SU sperm fraction, and 6.39 ± 2.18% (range: 1.58-10.37%) in BPG sperm fraction. Thus, the mean frequency of sperm cells with ssDNA fragmentation decreased: 3.14-fold (- 68.14%) in SU spermatozoa (p= 0.0317) and 3.29-fold (-69.6%) in BPG spermatozoa (p=0.0063) (Table 2 and Figure 4).

TUNEL assay evaluation revealed that the mean frequency of sperm with dsDNA and ssDNA fragmentation was 5.54 ± 2.53% (range: 3.00-9.00%) in non-fractionated spermatozoa, 5.5 ± 3.94% (range: 2.60-12.40%) in SU sperm fraction, and 2.76 ± 2.18% (range: 0.80-6.20%) in BPG sperm fraction. However, no significant differences between sperm fractions were found (Table 2 and Figure 4).

#### 5mC and 5hmC levels of sperm DNA

The mean frequency of sperm cells with highly methylated DNA (5mC) and hydroxymethylated DNA (5hmC), estimated for non-fractionated ejaculated sample and good-quality fractions (SU and BPG) was presented in Table 3.

**Table 3.**
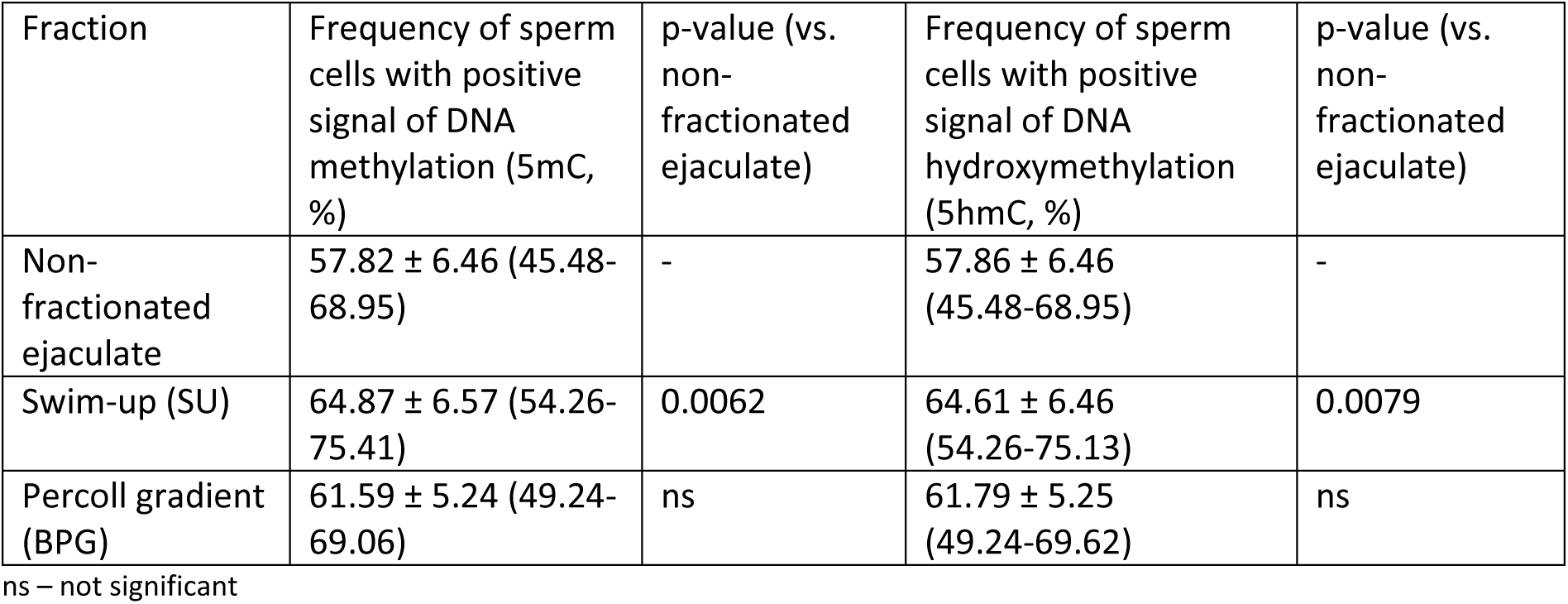
Mean global sperm DNA methylation (5mC) and hydroxymethylation (5hmC) status estimated for sperm cells from non-fractionated ejaculate and good-quality fractions (swim up fraction (SU) and Percoll gradient (BPG))

| Fraction | Frequency of sperm cells with positive signal of DNA methylation (5mC, %) | p-value (vs. non-fractionated ejaculate) | Frequency of sperm cells with positive signal of DNA hydroxymethylation (5hmC, %) | p-value (vs. non-fractionated ejaculate) |
| --- | --- | --- | --- | --- |
| Non-fractionated ejaculate | $57.82 \pm 6.46$ (45.48-68.95) | - | $57.86 \pm 6.46$ (45.48-68.95) | - |
| Swim-up (SU) | $64.87 \pm 6.57$ (54.26-75.41) | 0.0062 | $64.61 \pm 6.46$ (54.26-75.13) | 0.0079 |
| Percoll gradient (BPG) | $61.59 \pm 5.24$ (49.24-69.06) | ns | $61.79 \pm 5.25$ (49.24-69.62) | ns |

ns – not significant

The analysis revealed that at the levels of the mean 5mC or 5hmC frequencies were 57.82 ± 6.456% (45.48-68.95%) or 57.86 ± 6.46% (45.48-68.95%) (5mC or 5hmC, respectively) in non-fractionated ejaculate (Table 3). In contrast, the SU sperm fraction exhibited a significantly higher level of 5mC or 5hmC when compared to ejaculate, averaging: 64.87 ± 6.572% (54.26-75.41%) (p=0.0062) or 64.61 ± 6.456 (p=0.0079) (54.26%-75.13%), respectively (Table 3). On the other hand, no statistically significant differences were observed between BPG sperm fraction and ejaculate, where the average 5mC or 5hmC frequencies were 61.59 ± 5.241% (49.24-69.06%) or 61.79 ± 5.254% (49.24-69.62%), respectively (Table 3). Nonetheless, an increase of 6.52% and 6.79%, respectively, was observed in BPG sperm fraction vs. non-fractionated ejaculate.

### Sperm FISH analysis – chromosomal positioning

In this study we determined chromosomal positioning within the sperm cell nucleus through sequential stainings performed for the same individual sperm cell (cell by cell, *in situ* on a microscopic slide, as indicated in Figure 1, Figure 2 and Figure 3b). First, FISH was performed on spermatozoa of non-fractionated ejaculate and good-quality fractions (SU and BPG), followed by documentation of sperm cells’ position on microscopic slides. Second, immunofluorescence staining (IF) for 5mC and 5hmC was applied on the same slides, and subsequent topology measurements were performed. It allowed to establish the chromosomal positions in populations of sperm cells with low (hypo-) and high (hyper-) levels of DNA methylation/hydroxymethylation.

### Linear positioning

The linear positioning consisted of frequency values of each chromosome in three regions of the sperm nucleus and were presented in a Table 4 and in Figure 5 (mean values), supported by Additional file 3 (data for individual cases K1-K5).

**Figure 5.**
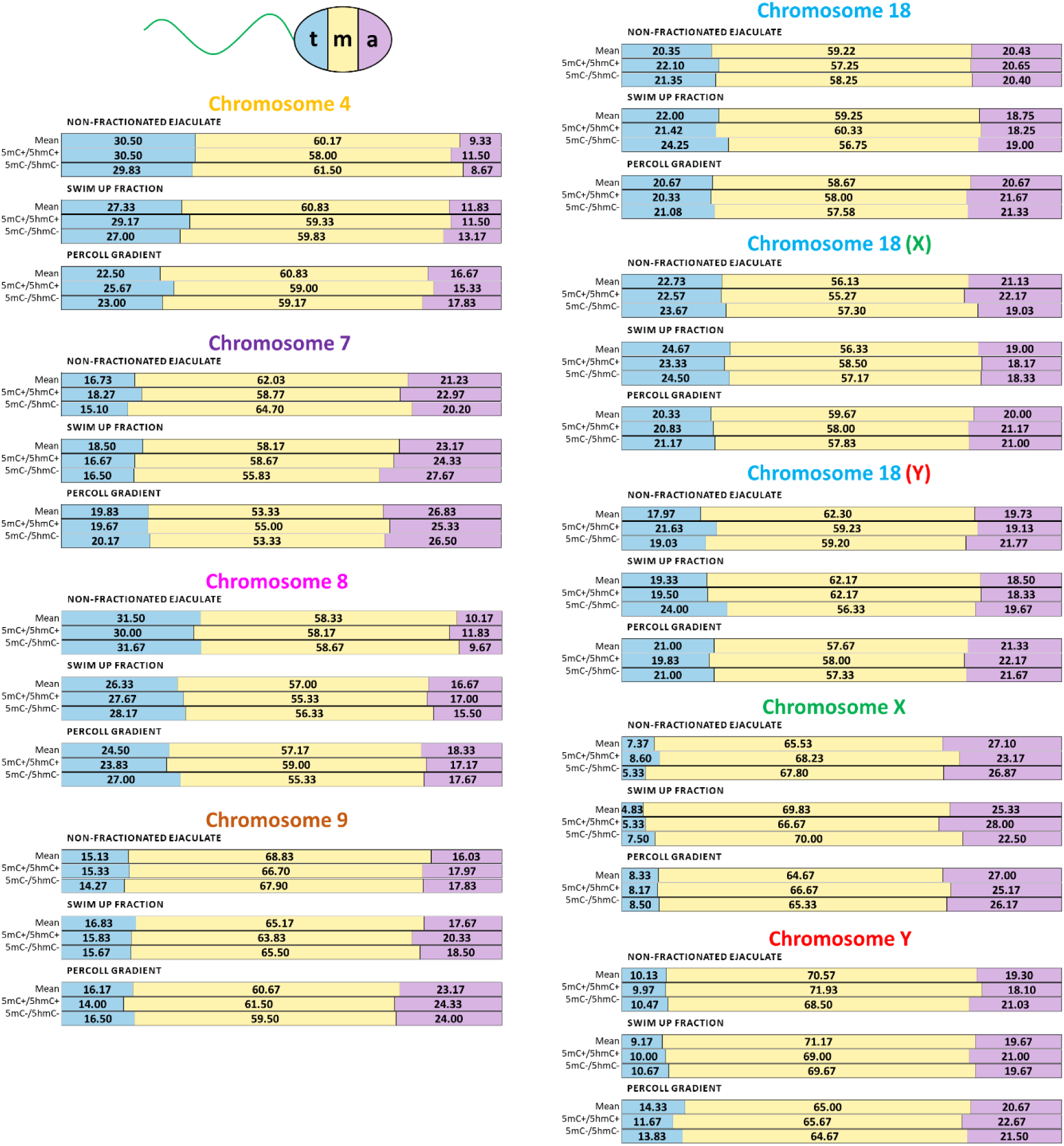
Linear positioning of centromeres within the sperm nucleus. Comparison of frequencies of chromosomes: 4, 7, 8, 9, 18, X, and Y in particular regions of the sperm cell nucleus (‘t’ – near the tail, ‘m’ – middle part, and ‘a’ – near the acrosome) in non-fractionated ejaculate, swim-up fraction (SU), and bottom layer of Percoll gradient (BPG), including also differentiation into high (5mC/5hmC+) and low methylated (5mC/5hmC-) sperm populations (according to the Table 4).

**Table 4.**
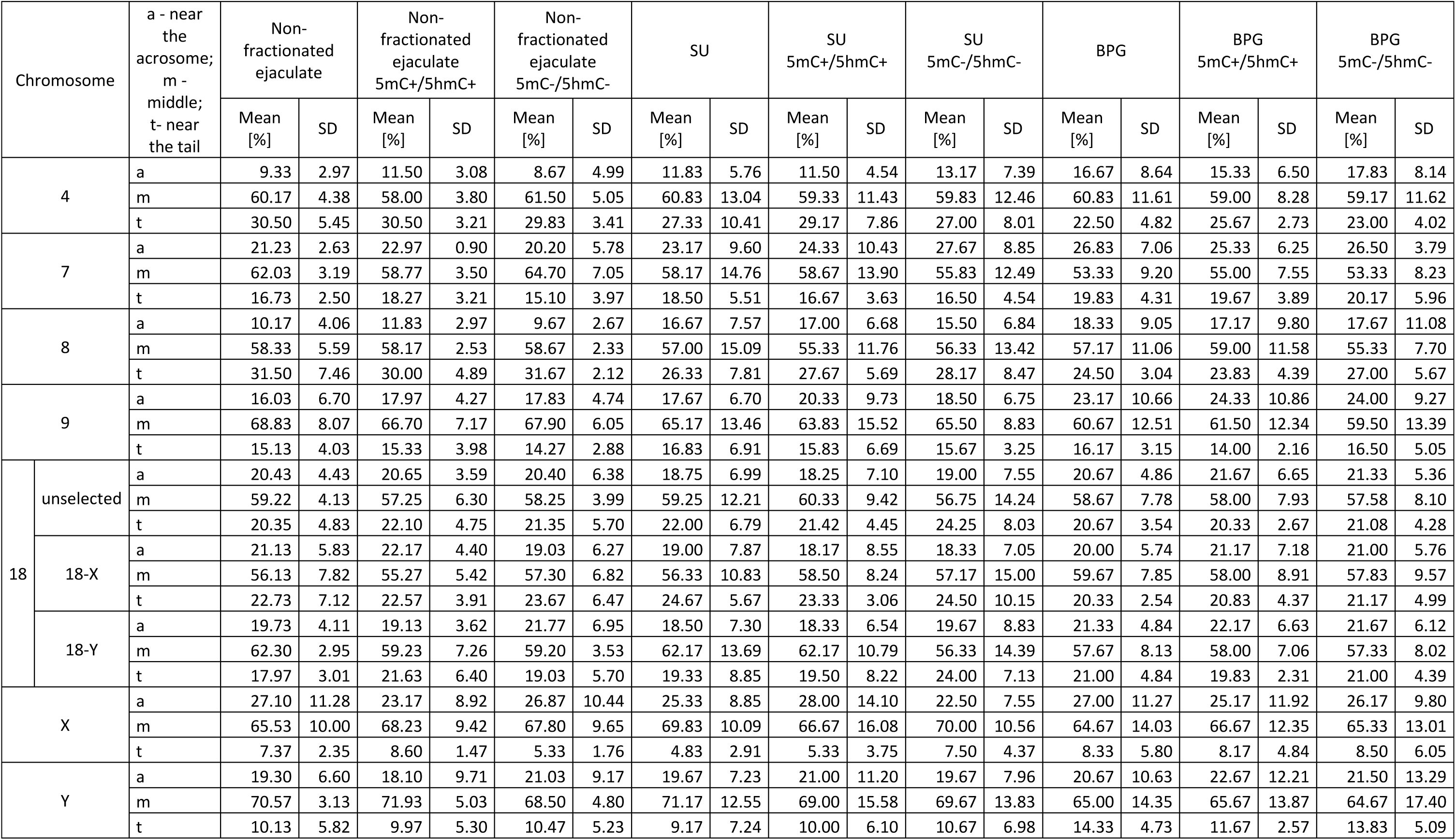
Linear positioning of centromeres of chromosomes: 4, 7, 8, 9, 18, X and Y in human spermatozoa of n=5 male normozoospermic cases (K1-K5). For each of the evaluated autosomes, the position of centromeres was defined for unselected spermatozoa according to sex chromosomes. For chromosome 18 additional measurements were done according to X- or Y-bearing spermatozoa. At least n=100 of signals were evaluated for each case and each chromosome. SU – swim up fraction, BPG – bottom layer of Percoll gradient, 5mC – global sperm DNA methylation, 5hmC – global sperm DNA hydroxymethylation.

Regardless of the fraction and the level of 5mC/5hmC, most chromosomes preferentially localized in the middle (‘m’) part of the sperm nucleus, with the exception of the X chromosome, which was predominantly found near the acrosome (‘a’), and chromosomes 4 and 8, mostly localised near the tail (‘t’). In fractions (SU and BPG), as well as in sperm with differentially methylated and hydroxymethylated DNA, there were no statistically significant differences in the localization of all evaluated chromosomes. However, the greatest difference in liner positioning was observed for chromosome 4, between non-fractionated ejaculate and BPG, where the centromeres were shifted towards the acrosomal area (‘a’: 9.33 ± 2.97% and 16.67 ± 8.64%, respectively; ‘m’: 60.17 ± 4.38% and 60.83 ± 11.61%, respectively; ‘t’: 30.50 ± 5.45% and 22.50 ± 4.82%, respectively) (Table 4).

### Non-fractionated ejaculate

All the evaluated chromosomes in non-fractionated ejaculate were preferentially localized in the middle part (‘m’) of the sperm nucleus (56.13-70.57%; mean: 62.57 ± 4.86%). Near the acrosome (‘a’), the frequency of centromeres varied from 9.33 to 27.10% (mean: 18.27 ± 5.62%), with the most acrosomal position noted for the centromere of chromosome X (27.10 ± 11.28%). Localization near the sperm tail (‘t’) ranged from 7.37 to 31.50% (mean: 19.16 ± 8.21%) and was preferentially exhibited by centromere of chromosome 4 (mean: 30.50 ± 5.45%) and 8 (31.50 ± 7.46%) (Table 4).

In non-fractionated ejaculate with highly methylated (5mC+) and hydroxymethylated (5hmC+) sperm DNA, all evaluated chromosomes were predominantly found in the middle part of the sperm nucleus (‘m’) ranging from 55.27 to 71.94% (mean: 61.51 ± 5.85%). The frequency of centromeres near the acrosome (‘a’) varied from 11.50 to 23.17% (mean: 18.61 ± 4.39%), with chromosome X exhibiting the most acrosomal localization (23.17 ± 8.92%). Centromeres positioned near the sperm tail (‘t’) were observed in 8.60 to 30.50% of cases (mean: 19.89 ± 7.74%), with chromosomes 4 and 8 showing the strongest preference for this localization (30.50 ± 3.21% and 30.00 ± 4.89%, respectively) (Table 4).

Similar observations have been revealed in non-fractionated ejaculate with low levels of methylated (5mC-) and hydroxymethylated (5hmC-) DNA. Linear positioning analysis revealed that the majority of evaluated chromosomes were preferentially located in the mid-region of the sperm nucleus (‘m’; 57.30-68.50%; mean: 62.65 ± 4.60%). Centromeres detected near the acrosomal region (‘a’) ranged from 8.67 to 26.87% (mean: 18.39 ± 5.80%), with chromosome X showing the most acrosomal localization (26.87 ± 10.44%). Centromeres found near the sperm tail (‘t’) were present in 5.33% to 31.67% of cases (mean: 18.97 ± 8.68%), with chromosomes 4 and 8 displaying the highest localization in this region (29.83 ± 3.41% and 31.67 ± 2.12%, respectively) (Table 4).

### Swim up fraction (SU)

In SU sperm fraction, the analysed chromosomes predominantly occupied the central part of the sperm nucleus (‘m’), with frequencies between 56.33% and 71.17% (mean: 62.21 ± 5.43%). Near the acrosomal region (‘a’), the centromere frequencies ranged from 11.83% to 25.33% (mean: 18.95 ± 3.82%), with chromosome X displaying the most acrosomal localization (25.33 ± 8.85%). Meanwhile, centromeres positioned close to the sperm tail (‘t’) accounted for 4.83% to 27.33% of cases (mean: 18.78 ± 7.63%), with chromosomes 4 and 8 being the most frequently localized in this area (27.33 ± 10.41% and 26.33 ± 7.81%, respectively) (Table 4).

In SU sperm fraction with high levels of 5mC+ and 5hmC+ sperm chromosomes were primarily distributed in the midsection of the sperm nucleus (‘m’), with frequencies varying from 55.33% to 69.00% (mean: 61.54 ± 4.33%). Centromeres detected in the acrosome (‘a’) area ranged between 11.50% and 28.00% (mean: 19.66 ± 4.65%), with chromosome X exhibiting the most acrosomal positioning (28.00 ± 14.10%). Meanwhile, centromeres located towards the sperm tail (‘t’) were observed in 5.33% to 29.17% of cases (mean: 18.77 ± 7.80%), with chromosomes 4 and 8 showing the highest association for this region (29.17 ± 7.86% and 27.67 ± 5.69%, respectively) (Table 4).

In SU sperm fraction with 5mC- and 5hmC- low levels there was a preferential localization of the chromosomes in the middle section of the sperm nucleus (‘m’) revealed, with frequencies ranging from 55.83% to 70.00% (mean: 60.82 ± 5.92%). The centromeres located near the acrosomal region (‘a’) varied between 13.17% and 27.67% (mean: 19.33 ± 4.10%), with chromosome 7 being the most acrosomally positioned (27.67 ± 8.85%). Conversely, centromeres close to the sperm tail (‘t’) were found in 7.50% to 28.17% of cases (mean: 19.81 ± 7.45%), with the highest preference for this region of chromosomes 4 and 8 (27.00 ± 8.01% and 28.17 ± 8.47%, respectively) (Table 4).

### Bottom layer of Percoll gradient (BPG)

In BPG sperm fraction most of analysed chromosomes were found in the midsection of the sperm nucleus (‘m’), with values ranging from 53.33% to 65.00% (mean: 59.74 ± 3.66%). Centromeres positioned near the acrosomal region (‘a’) were detected in 16.66% to 27.00% of cases (mean: 21.63 ± 3.51%), with chromosome X being the most acrosomally localized (27.00 ± 11.27%). Meanwhile, centromeres located near the sperm tail (‘t’) varied between 8.33 and 24.50% (mean: 18.63 ± 4.92%), with chromosomes 4 and 8 predominantly found in this area (22.50 ± 4.82% and 24.50 ± 3.04%, respectively) (Table 4).

In BPG sperm fraction with high 5mC+ and 5hmC+ levels, it was demonstrated that the majority of examined chromosomes were predominantly situated also in the mid-region of the sperm nucleus (‘m’), with frequencies between 55.00% and 66.67% (mean: 60.09 ± 3.84%). Centromeres positioned near the acrosomal area (‘a’) were observed in 15.33% to 25.33% of cases (mean: 21.67 ± 3.44%), with chromosome 7 showing the highest acrosomal presence (25.33 ± 6.25%). Meanwhile, centromeres localized near the sperm tail (‘t’) exhibited frequencies from 8.17% to 25.67% (mean: 18.22 ± 5.75%), with chromosomes 4 and 8 being the most frequently observed in this region (25.67 ± 2.73% and 23.83 ± 4.39%, respectively) (Table 4).

Similarly, the distribution of chromosomes in BPG sperm fraction with 5mC- and 5hmC- low level showed that the majority of them were concentrated in the mid-region of the sperm nucleus (‘m’), with frequencies spanning from 53.33% to 65.33% (mean: 58.90 ± 3.94%). Centromeres near the acrosomal region (‘a’) were found in 17.67% to 26.50% of cases (mean: 21.96 ± 8.07%), with chromosome 7 occupying the most acrosomal position (26.50 ± 3.79%). Meanwhile, centromeres located towards the sperm tail (‘t’) were detected in 8.50% to 27.00% of cases (mean: 19.14 ± 5.45%), with chromosomes 4 and 8 showing a preferential localization for this region (23.00 ± 4.02% and 27.00 ± 5.67%, respectively) (Table 4).

### Radial positioning

The radial positioning results were presented in Tables: 5 and 6, Figure 6, and Additional files: 4, 5 and 6.

**Figure 6.**
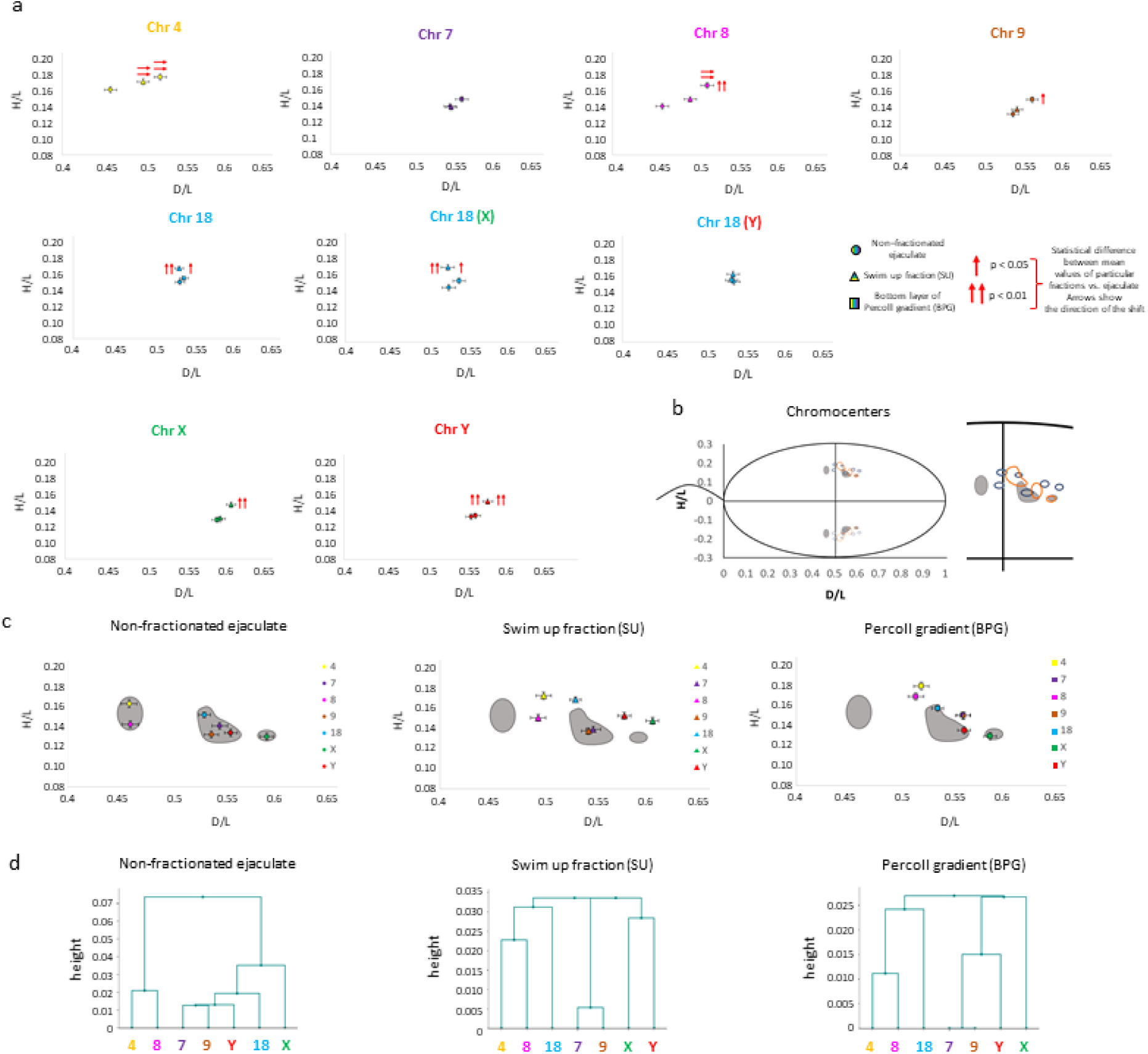
Radial positioning of the centromeres of examined chromosomes (4, 7, 8, 9, 18, X, Y) within the human sperm cell nucleus. Three populations of spermatozoa were evaluated in n=5 normozoospermic cases (K1-K5) in: non-fractionated ejaculate, swim up fraction (SU) and Percoll gradient (BPG). Positioning of the centromeres was determined as the points on coordinate system: D/L ± SE (OX axis) and H/L ± SE (OY axis) (acc. to data in Table 5). **a** Positioning results for individual centromeres (circle: non-fractionated ejaculate, triangle: swim up fraction (SU), square: bottom layer of Percoll gradient (BPG)). Localizations that differ significantly from the non-fractionated ejaculate were indicated by arrows: double arrow for p < 0.01, single arrow for p < 0.05. Arrows also indicate the direction of the observed shift/repositioning of a centromere. Bars show standard errors (SE). **b** Schematic region within the sperm nucleus occupied by chromocenters (solid lines), along with the investigated centromeres. Dotted lines represent their mirror images; non-fractionated ejaculate (grey areas), swim- up fraction (SU; dark blue), and Percoll gradient (BPG; orange). **c** The chromocenter areas represented for each sperm population, with the grey colour for non-fractionated ejaculate as a background for visualization of shifts when compared to swim up (SU) and Percoll gradient (BPG) fractions. **d** Positions of analysed centromeres represented by hierarchical Ward clustering analysis showing the reciprocal positions of selected chromosomal pairs.

**Table 5.** Radial positioning of centromeres of chromosomes: 4, 7, 8, 9, 18, X and Y in human spermatozoa of n=5 male normozoospermic cases (K1-K5). For each of the evaluated autosomes, the position of centromeres was defined for unselected spermatozoa according to sex chromosomes. For chromosome 18 additional measurements were done according to X- or Y-bearing spermatozoa. At least n=100 of signals were evaluated for each case and each chromosome. SU – swim up fraction, BPG – bottom layer of Percoll gradient, 5mC – global sperm DNA methylation, 5hmC – global sperm DNA hydroxymethylation, D/L – signal distance along the tail-acrosome axis, H/L – signal proximity to the nuclear periphery (acc. to Fig. 2), SE – standard error of the mean.

| Chromosome |  | Non-fractionated ejaculate | Non-fractionated ejaculate 5mC+/5hmC+ | Non-fractionated ejaculate 5mC-/5hmC- | SU | SU 5mC+/5hmC+ | SU 5mC-/5hmC- | BPG | BPG 5mC+/5hmC+ | BPG 5mC-/5hmC- |
| --- | --- | --- | --- | --- | --- | --- | --- | --- | --- | --- |
| 4 | D/L | 0.462 | 0.465 | 0.465 | 0.504<sup>→→</sup> | 0.495 | 0.503<sup>→→</sup> | 0.525<sup>→→</sup> | 0.513<sup>→→</sup> | 0.517<sup>→→</sup> |
|  | SE | 0.008 | 0.008 | 0.008 | 0.007 | 0.008 | 0.008 | 0.008 | 0.008 | 0.008 |
|  | H/L | 0.163 | 0.162 | 0.163 | 0.173 | 0.173 | 0.170 | 0.179 | 0.176 | 0.169 |
|  | SE | 0.004 | 0.004 | 0.004 | 0.004 | 0.004 | 0.004 | 0.004 | 0.004 | 0.004 |
| 7 | D/L | 0.552 | 0.556 | 0.558 | 0.553 | 0.561 | 0.567 | 0.567 | 0.566 | 0.562 |
|  | SE | 0.008 | 0.008 | 0.008 | 0.007 | 0.007 | 0.007 | 0.008 | 0.008 | 0.008 |
|  | H/L | 0.141 | 0.149 | 0.141 | 0.140 | 0.143 | 0.144 | 0.150 | 0.150 | 0.149 |
|  | SE | 0.004 | 0.004 | 0.004 | 0.003 | 0.003 | 0.003 | 0.004 | 0.004 | 0.004 |
| 8 | D/L | 0.463 | 0.479 | 0.464 | 0.498 | 0.497 | 0.493 | 0.520<sup>→→</sup> | 0.508<sup>→→</sup> | 0.507<sup>→→</sup> |
|  | SE | 0.008 | 0.008 | 0.008 | 0.008 | 0.008 | 0.008 | 0.008 | 0.008 | 0.008 |
|  | H/L | 0.142 | 0.156 | 0.136 | 0.151 | 0.150 | 0.159<sup>↑</sup> | 0.169<sup>↑↑</sup> | 0.162<sup>↑↑</sup> | 0.157 |
|  | SE | 0.004 | 0.004 | 0.004 | 0.003 | 0.004 | 0.004 | 0.004 | 0.004 | 0.004 |
| 9 | D/L | 0.543 | 0.552 | 0.550 | 0.548 | 0.556 | 0.548 | 0.567 | 0.574 | 0.570 |
|  | SE | 0.007 | 0.007 | 0.007 | 0.007 | 0.007 | 0.007 | 0.007 | 0.007 | 0.007 |
|  | H/L | 0.132 | 0.143 | 0.136 | 0.138 | 0.144 | 0.139 | 0.150<sup>↑</sup> | 0.149<sup>↑</sup> | 0.146 |
|  | SE | 0.004 | 0.004 | 0.003 | 0.003 | 0.003 | 0.003 | 0.004 | 0.004 | 0.004 |

| Chromosome |  |  | Non-fractionated ejaculate | Non-fractionated ejaculate<br>5mC+/5hmC+ | Non-fractionated ejaculate<br>5mC-/5hmC- | SU | SU<br>5mC+/5hmC+ | SU<br>5mC-/5hmC- | BPG | BPG<br>5mC+/5hmC+ | BPG<br>5mC-/5hmC- |
| --- | --- | --- | --- | --- | --- | --- | --- | --- | --- | --- | --- |
| 18 | unselected | D/L | 0.536 | 0.533 | 0.537 | 0.535 | 0.532 | 0.536 | 0.541 | 0.545 | 0.539 |
|  |  | SE | 0.006 | 0.006 | 0.006 | 0.006 | 0.005 | 0.006 | 0.005 | 0.006 | 0.005 |
|  |  | H/L | 0.152 | 0.151 | 0.151 | 0.169<sup>↑↑</sup> | 0.170<sup>↑↑</sup> | 0.169<sup>↑↑</sup> | 0.157<sup>a</sup> | 0.156 | 0.160 |
|  |  | SE | 0.003 | 0.003 | 0.003 | 0.003 | 0.002 | 0.003 | 0.002 | 0.002 | 0.003 |
|  | 18-X | D/L | 0.531 | 0.536 | 0.530 | 0.530 | 0.525 | 0.536 | 0.544 | 0.547 | 0.542 |
|  |  | SE | 0.008 | 0.009 | 0.008 | 0.008 | 0.008 | 0.008 | 0.008 | 0.008 | 0.008 |
|  |  | H/L | 0.148 | 0.155 | 0.149 | 0.173<sup>↑↑</sup> | 0.172<sup>↑↑</sup> | 0.171<sup>↑↑</sup> | 0.156<sup>a</sup> | 0.157 | 0.160 |
|  |  | SE | 0.004 | 0.004 | 0.004 | 0.004 | 0.003 | 0.004 | 0.004 | 0.003 | 0.004 |
|  | 18-Y | D/L | 0.542 | 0.530 | 0.544 | 0.541 | 0.539 | 0.535 | 0.539 | 0.544 | 0.535 |
|  |  | SE | 0.008 | 0.008 | 0.008 | 0.008 | 0.008 | 0.008 | 0.008 | 0.008 | 0.008 |
|  |  | H/L | 0.156 | 0.147 | 0.153 | 0.165 | 0.169 | 0.167 | 0.157 | 0.154 | 0.160 |
|  |  | SE | 0.004 | 0.004 | 0.004 | 0.004 | 0.004 | 0.004 | 0.003 | 0.003 | 0.004 |
| X |  | D/L | 0.598 | 0.585 | 0.605 | 0.612 | 0.617 | 0.606 | 0.594 | 0.595 | 0.592 |
|  |  | SE | 0.007 | 0.007 | 0.006 | 0.006 | 0.006 | 0.006 | 0.006 | 0.006 | 0.006 |
|  |  | H/L | 0.130 | 0.124 | 0.134 | 0.148<sup>↑↑</sup> | 0.147<sup>↑↑</sup> | 0.153<sup>↑↑</sup> | 0.129 | 0.133 | 0.134 |
|  |  | SE | 0.004 | 0.003 | 0.004 | 0.003 | 0.003 | 0.003 | 0.003 | 0.003 | 0.003 |
| Y |  | D/L | 0.563 | 0.562 | 0.573 | 0.584 | 0.587 | 0.577 | 0.568 | 0.573 | 0.565 |
|  |  | SE | 0.006 | 0.006 | 0.007 | 0.006 | 0.006 | 0.007 | 0.007 | 0.006 | 0.007 |
|  |  | H/L | 0.134 | 0.132 | 0.136 | 0.153<sup>↑↑</sup> | 0.154<sup>↑↑</sup> | 0.148 | 0.135<sup>b</sup> | 0.142 | 0.134<sup>b</sup> |
|  |  | SE | 0.004 | 0.004 | 0.004 | 0.003 | 0.003 | 0.003 | 0.003 | 0.003 | 0.003 |

Mean values statistically significant ( $p < 0.05$ ) from the mean values of non-fractionated ejaculate sperm are marked with arrows:
↑ centromere localized closer to the periphery of the nucleus
↓ centromere localized closer to the centre of the nucleus
→ centromere localized closer to the acrosome
← centromere localized closer to the tail.
Mean values with high statistical significance ( $p < 0.01$ ) from the mean values of non-fractionated ejaculate sperm are marked with double arrows:
↑↑ centromere localized closer to the periphery of the nucleus
↓↓ centromere localized closer to the centre of the nucleus
→→ centromere localized closer to the acrosome

The spatial positioning of all the studied centromeres in non-fractionated ejaculate as well as in both good quality sperm fractions (SU and BPG) was limited to a restricted small area located in the central part of the nucleus, towards its apical side (Fig. 6a,b). Analysis revealed also several changes in the chromosome order within the sperm nucleus. The detailed results were presented in Table 5 and 6 and Fig. 6a.

The results based on the ’tail–acrosome’ criterion consistently demonstrated that, across all the analysed fractions, centromeres of chromosomes X and Y were predominantly located in the acrosomal region, while centromeres of chromosomes 4 and 8 were positioned closest to the sperm tail. Notably, the apical localization of centromere of chromosome X was observed in all analysed samples (Fig. 6c). In SU sperm fraction, centromere of chromosome X was repositioned towards peripheral area, when compared to non-fractionated spermatozoa (p=0.0047) and BPG sperm fraction (p=0.0021) (Fig. 6a). Centromere of chromosome Y in SU sperm fraction, moved towards peripheral area, when compared to non-fractionated spermatozoa (p=0.0011) and BPG (p=0.0071) (Fig. 6a).

According to the ‘centre–periphery’ criterion, chromosome 4 consistently occupied the most peripheral position across all the analysed fractions. In contrast, in the non-fractionated ejaculate and BPG sperm fraction, chromosomes X and Y were located most centrally, deep within the sperm nucleus, whereas in the SU sperm fraction, chromosomes 7 and 9 assumed the most central positions (Fig. 6a and Table 5). Furthermore, centromere of chromosome 18 shifted towards peripheral area in SU sperm fraction, when compared to non-fractionated spermatozoa (p<0.0001) and BPG sperm fraction (p=0.0467) (Fig. 6a). When considering positioning of centromere of chromosome 18 in sperm cells carrying chromosome X centromere, chromosome 18 centromere shifted towards peripheral area in SU fracction, when compared to non-fractionated spermatozoa (p<0.0001) and BPG sperm fraction (p=0.036) (Fig. 6a). When considering positioning of centromere of chromosome 18 in sperm cells carrying chromosome Y potential shift of centromere of chromosome 18 was not statistically significant (p>0.05) (Fig. 6a).

For SU and BPG sperm fractions the chromosome 4 was repositioned towards the acrosomal area of sperm cell nucleus when compared to non-fractionated sperm (p=0.0079 and p<0.0001, respectively) (Fig 6d). There was no statistically significant change in positioning of centromere of chromosome 7 (p>0.05) (Fig 6d). Chromosome 8 in BPG sperm fraction shifted towards the acrosome area (p<0.0001) or peripheral area (p<0.0001) of the sperm cell nucleus when compared to non-fractionated sperm (Fig. 6d). It was also observed that chromosomes 4 and 8 colocalized and assumed a position in the middle part of sperm nucleus in non-fractionated ejaculate, SU and BPG sperm fractions (Fig 6a, c) and also in each individual case (Additional file 6). Chromosome 9 shifted towards peripheral area in BPG sperm fraction when compared non-fractionated sperm (p=0.0118) (Fig. 6a). Summing up, all analysed chromosomes exhibited some changes in their localization within the good-quality fractions, except for chromosome 7, which retained its stable position across all the fractions (Table 6).

**Table 6.** Summary of repositioned centromeres among evaluated chromosomes between fractions. SU – swim up fraction, BPG – bottom layer of Percoll gradient. ‘Tail-acrosome’ means that centromere was shifted from the tail to the acrosomal region. ‘Centre-periphery’ means the repositioning towards the nuclear periphery. ‘Both’ stands for centromere repositioning observed in both directions: tail-acrosome and centre-periphery, while ‘None’ for no statistically significant change in centromere localization between fractions.

|  | Direction of repositioning |  |  |  |
| --- | --- | --- | --- | --- |
|  | Tail-acrosome | Centre-periphery | Both | None |
| SU vs. Non-fractionated ejaculate | 4 | 18 |  | 7 |
|  |  | X |  | 8 |
|  |  | Y |  | 9 |
| BPG vs. Non-fractionated ejaculate | 4 | 9 | 8 | 7 |
|  |  |  |  | 18 |
| SU vs. BPG |  | 18 |  | 4 |
|  |  | X |  | 7 |
|  |  | Y |  | 8 |
|  |  |  |  | 9 |

While evaluating the positioning in differentially methylated (5mC) and hydroxymethylated (5hmC) sperm cells, there were no significant differences in the positioning of all analysed chromosomes within the same fraction (non-fractionated ejaculate, SU and BPG) (p > 0.05) (Table 5).

Hierarchical Ward clustering showed that in non-fractionated ejaculated sample analysed chromosomes were localized pair-wise (4 and 8), in a group (7, 9, 18 and Y) or remained single (X) (Fig 6d). In non-fractionated ejaculate three chromocenter areas were visible (Fig. 6c). In comparison, in SU sperm fraction chromosomes were localized pair-wise (4 and 8, 7 and 9, X and Y) or remained single (18), and in BPG sperm fraction chromosomes were localized in two group (4, 8, 18 and 7, 9, Y, X) (Fig 6d). In SU and BPG sperm fractions chromocenter occupied smaller area than in non-fractionated sperm (Fig. 6b, c, d). This suggests the presence of distinct chromocenter fragment in each sperm fraction and highlights the differences in chromosomal distribution.

### Distances between the chromosomes’ centromeres

The average distances between centromeres of selected chromosome pairs (4 vs. 8, 7 vs. 9, 18 vs. X, 18 vs. Y) were analysed across the examined sperm fractions. The results of evaluation were presented in the Table 7, Figure 7 and Additional file 6. In line with previous reports of possible interindividual variability in localization of chromosomes (23,105,106,112), also in this study we have examined this aspect (Additional file 5).

**Figure 7.**
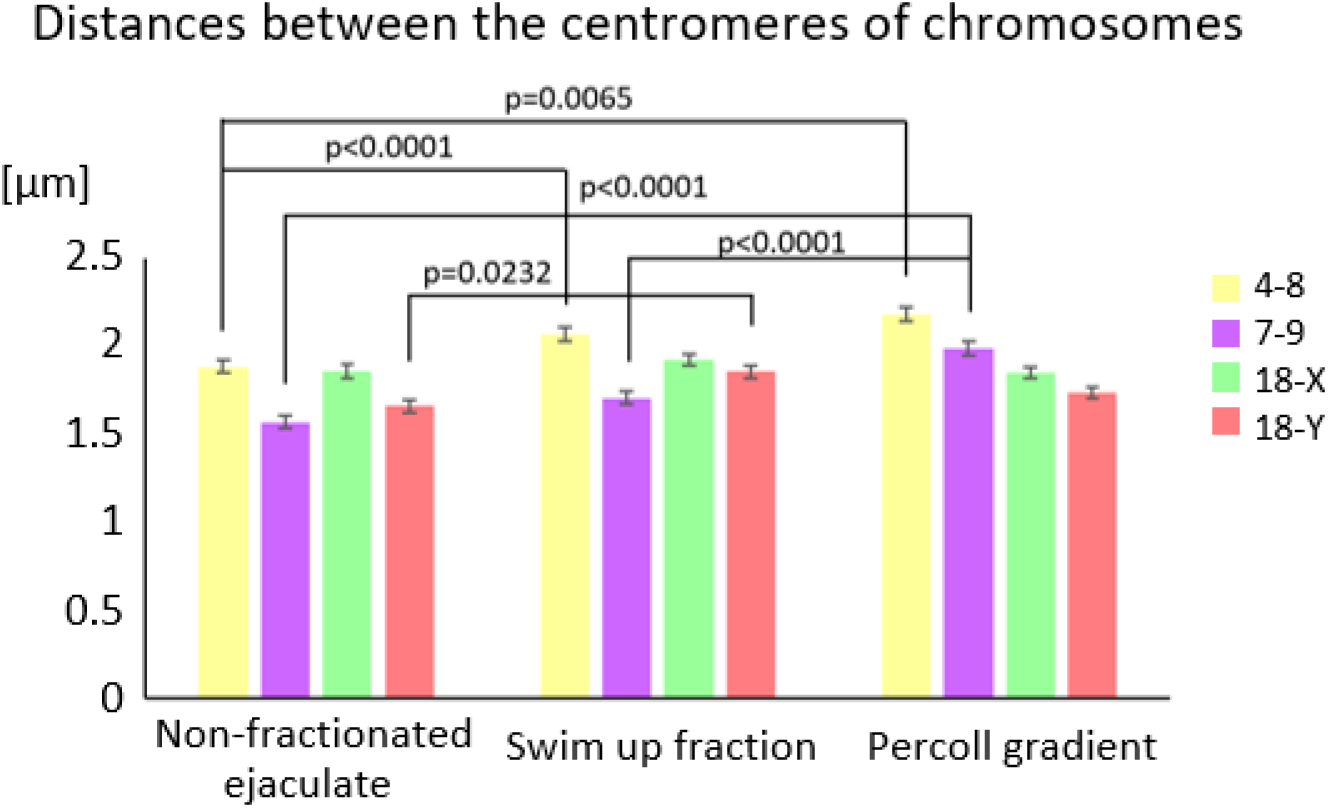
Distances between the centromeres of chromosomes 4 vs. 8, 7 vs. 9, 18 vs. X, 18 vs. Y in non-fractionated ejaculate, swim up fraction (SU) and Percoll gradient (BPG) in n=5 normozoospermic cases (K1-K5). Statistical significance was considered at p < 0.05.

**Table 7.** Distances between the centromeres of chromosomes 4 and 8, 7 and 9, 18 and X, 18 and Y in non-fractionated ejaculate and good-quality fractions (SU and BPG).

| The average distance between chromosomes [$\mu\text{m}$] | | | | |
| --- | --- | --- | --- | --- |
| Chromosomes |  | Non-fractionated ejaculate | SU | BPG |
| 4-8 | Mean | 1.888 | 2.073** | 2.184** |
|  | SE | 0.03797 | 0.03953 | 0.04091 |
| 7-9 | Mean | 1.571 | 1.707 | 1.989**# |
|  | SE | 0.034 | 0.03674 | 0.03991 |
| 18-X | Mean | 1.859 | 1.926 | 1.853 |
|  | SE | 0.04028 | 0.03576 | 0.03439 |
| 18-Y | Mean | 1.662 | 1.858* | 1.739 |
|  | SE | 0.03512 | 0.03596 | 0.03566 |

\* values different significantly from the mean value of non-fractionated sperm ( $p < 0.05$ )
\*\* values different significantly from the mean value of non-fractionated sperm ( $p < 0.01$ )

There were no significant differences in distances between spermatozoa with high or low 5mC and 5hmC levels (p>0.05), independently of the analysed sperm fraction.

In both SU and BPG sperm fractions, the distance between the centromeres of chromosomes 4 and 8 was wider when compared to non-fractionated spermatozoa, by 1.1-fold in SU sperm fraction (p = 0.0065) and by 1.16-fold in BPG sperm fraction (p<0.0001) (Fig. 7). Chromosomes 4 and 8 consistently colocalized in all three analysed sperm populations and the interindividual variability remained at the negligible level (with 0 out of 5 variable cases to 2 out of 5 noted in SU or BPG sperm fractions) (Additional file 5).

When considering the distance between centromeres of chromosomes 7 and 9, it was found that in BPG sperm fraction this value was 1.27-fold higher than in non-fractionated spermatozoa (p<0.0001), and 1.17-fold higher when compared to SU sperm fraction (p<0.0001) (Fig. 7). Chromosomes 7 and 9 were located similarly: each showed variability in 1 out of 5 cases in native ejaculated sample , 3 out of 5 or 4 out of 5 in SU sperm fraction, respectively, and 2 out of 5 in BPG sperm fraction. Despite high variability in SU sperm fraction, the colocalization of chromosomes 7 and 9 remained stable (Additional file 5, Fig 6a).

A distance between centromeres of chromosomes 18 and Y was 1.12-fold higher in SU sperm fraction than in non-fractionated ejaculate (p=0.0232) (Fig. 7). For the Y chromosome, we have documented that both fractioning methods had a positive influence to stabilize Y chromosome position (from 3 out of 5 cases in native ejaculated sample, 2 out of 5 in SU sperm fraction, to 1 out of 5 in BPG sperm fraction) (Additional file 5). In Y bearing spermatozoa, chromosome 18 displayed almost no variability, what seems to underline the independent positioning pattern of those chromosomes, confirmed also by the changed distances between them, depending from the observed sperm fraction (1.12-fold higher in SU sperm fraction than in non-fractionated ejaculate (p=0.0232) (Fig. 7; Additional file 5)).

In case of X-bearing spermatozoa, the variability of chromosome 18 position increased from none in native ejaculated sample, to 1 out of 5 cases in SU sperm fraction, and 2 out of 5 in BPG sperm fraction, while the variability of chromosome X positioning remained on a middle level (3 out of 5 cases in native ejaculated sample and SU sperm fraction and 2 out of 5 in BPG sperm fraction) (Additional file 5). This observation may suggest some kind of linkage between reciprocal positioning of chromosome 18 and X, what can be supported by the fact of unchanged distances between them (p>0.05)(Fig 7).

## Discussion

This is the first study focusing on differences in chromosomal positioning within sperm cell nucleus in high-quality spermatozoa selected *via* swim-up or centrifugation in a density gradient, techniques used in infertility clinics for *in vitro* fertilization. Sequential staining algorithm was applied to analyse both: chromosomal localization, as well as the level of epigenetic modifications – specifically: DNA methylation (5mC) and hydroxymethylation (5hmC) – on the same individual sperm cell. The experimental algorithm was conducted step by step and involved: (i) the separation of high-quality fractions, (ii) topological measurements, and (iii) the assessment of global levels of sperm DNA 5mC and 5hmC, with already determined positioning of evaluated chromosomes. This integrated approach allowed us to correlate the spatial organization of chromosomes with the epigenetic profile of spermatozoa across different sperm quality fractions. By combining all molecular data obtained, including also characteristics of sperm chromatin integrity (chromatin protamination and DNA fragmentation), the study also focused on sperm quality indicators, in the paternal chromosomal context.

Initially, we performed a qualitative assessment of sperm chromatin to determine the structural integrity and overall quality of the sperm DNA in evaluated males. Our findings revealed that the frequency of high-quality spermatozoa with properly protaminated chromatin was significantly higher in selected good quality sperm fractions compared to non-fractionated native ejaculated samples. This finding aligns with previous studies demonstrating reduced histone retention in gradient-selected sperm fractions (69). It was also shown that increased chromatin condensation may protect critical regions of the sperm genome, including genes involved in early embryonal development. Regions retaining histones may also harbour genes necessary for these early development processes (113,114). It has been documented that imbalance in the ratio of protamines to the remaining histones, has been implicated in male infertility (22–24), revealed reduction of sperm quality or induction of sperm DNA damage (115–117), ultimately leading to breakdown of the fertilization process or decreased fertilization rate (118) and/or embryonic development (68,85,119). It was also shown, that poor sperm chromatin protamination negatively affects fertilization rates in ICSI (intracytoplasmic sperm injection) procedure involving gametes from healthy donors (113). These data followed by findings in our study emphasize the importance of chromatin integrity and proper DNA protamination for successful fertilization and embryogenesis and highlight the need for careful sperm selection.

In our study, no significant changes were observed in DNA fragmentation levels across the analysed intact and fractionated sperm samples using the TUNEL assay, which detects both single-stranded (ssDNA) and double-stranded DNA (dsDNA) breaks. This result may be attributed to the fact that the study population exclusively consisted of control men without a history of reproductive failures, thus, demonstrated low baseline levels of DNA damage seem to be reasonable. On the other hand, in previous studies performed with infertile patient groups the altered levels of ssDNA and dsDNA breaks were clearly documented (69,120). What is interesting and still widely discussed, there are ongoing discrepancies concerning methods used for sperm DNA fragmentation, supported by data originated from different laboratories. However, it has been shown that in motile sperm population selected by density-gradient centrifugation, from infertile patients with normal and abnormal spermiograms, there was a significant decrease in the frequency of sperm with damaged DNA observed (121,122), whereas in the applied swim-up method the recovered sperm showed no significant improvement (123). In contrast, other studies found that the frequency of sperm with fragmented DNA was reduced significantly in swim up-treated but not after density-gradient centrifugation, compared with the whole semen in infertile normozoospermic or oligozoospermic patients (124). In our study, we also applied the acridine orange test (AO), which specifically detects single-stranded DNA breaks. We have documented statistically significant differences between swim-up (SU) and Percoll gradient (BPG) spermatozoa which exhibited decreased level of ssDNA fragmentation when compared to the non-fractionated ejaculate. These observations may indicate that the efficacy of sperm DNA fragmentation reduction may vary, being dependent on the type of assay used (relationship to particular type of DNA damage), and studied male population, highlighting the need for standardization of approaches used for this purpose. During IVF procedures, especially ICSI, sperm selection from the whole non-fractionated ejaculate, has been rather a random process that may result in an unintended use of DNA-damaged spermatozoa – DNA fragmentation is not visible microscopically during the selection for IVF and thus, sperm cells morphologically correct can carry on highly fragmented sperm DNA (98). Therefore, it is important to assess the degree of DNA fragmentation in human sperm, as this parameter has been linked to fertilization outcomes, embryo development, pregnancy success and reduction of the risk of transmitting genetic damage to offspring (69,98,98,120–127). What is also crucial is the fact that spermatozoa lack of the repair mechanisms and if the level of a sperm DNA damage exceeds the oocyte’s capacity to repair it, then the cells might undergo apoptosis (26,128).

The analysis of global levels of DNA methylation (5mC) and hydroxymethylation (5hmC) revealed high levels of both epimarks in the good-quality sperm fractions compared to the non-fractionated ejaculate. A statistically significant difference was observed exclusively in SU sperm fraction, in which the levels were approximately 12% higher than in the native ejaculated samples. In BPG sperm fraction, 5mC and 5hmC levels were still approximately 6.5% higher, indicating a positive potential trend, however without the statistical significance. In previous studies it was shown that spermatozoa from fractionated samples may reflect different methylation patterns (69,129,130). Similarly, in fertile normozoospermic males the 5mC values observed were higher than in infertile patients (101,131). Changes in gene methylation pattern were also well documented in normozoospermic but infertile males (132,133). Previous reports indicated that the pregnancy rates were higher when sperm DNA methylation was above a certain threshold (134,135). Thus, it can be suggested that improved sperm selection techniques may be able to select a sperm population with proper epigenetic profile. Future studies need to determine what is considered normal in epigenetics and establish the real risks important for embryo development and the health of offspring (129). It has to emphasized is important to note that our study focused on global DNA methylation and hydroxymethylation in a group of normozoospermic men without known fertility problems, thus, the lack of significant differences may be due to the fact that this group was quite homogeneous. More spread out differences in methylation pattern might be found in more diverse populations, especially those including men with impaired spermatogenesis or reproductive failures (101,102).

Nuclear sperm architecture has been supposed to influence the early embryonic development (136,137), including the later stages of zygote development (138). It is suggested that abnormal chromosomal positioning within the sperm nucleus can have a negative impact on fertilisation and early embryogenesis (139), highlighting the importance of chromatin organization in reproductive success. Changes in sex chromosome positioning have been reported in semen samples with impaired sperm parameters (140), while abnormal nuclear organization has been associated with defective sperm head morphology (141).

Disturbances in the nuclear positioning of the chromocenters may be linked to compromised spermatogenesis, suggesting that the structural integrity and position of chromosomal territories are crucial for normal sperm function (127). In spermatozoa, centromeres of non-homologous chromosomes have a tendency to aggregate when composing chromocenters (142–144). It was shown that, in immotile spermatozoa chromocenter formation was disrupted (143). In previous studies it was demonstrated that less than 10% of spermatozoa contained a single chromocenter, with a common tendency to be located in the central region (139,140,143). There were also data revealing the existence of 1-3 chromocenters in 73% of cells (143). Our findings confirmed previously published data that centromeres of chromosomes have a defined and stable position in non-fractionated spermatozoa from native ejaculated samples (12,23,106). In our study the number of chromocenters varied between 2 and 4. A more fragmented arrangement of chromocenters may play a key role in the ordered release of paternal chromosomes after fertilization (8). Ioannou et al. suggested that specific chromosomes may preferentially contribute to the formation of individual chromocenters (8). In our study, the reduction of chromocenter area was observed, which is not surprising when our data were focused on selected good-quality sperm fractions (SU, BPG), which is presented for the first time. The link should be pointed out between the sperm quality (also at the chromosomal organization level) and previously documented wider area of chromocenters in males with disrupted spermatogenesis (103,105,106,143).

In our study, chromosome 4 relocated towards the acrosome in both good-quality sperm fractions, while chromosomes 18, X, and Y shifted to the nuclear periphery in motile sperm. Additionally, in sperm with good motility and morphology, chromosome 8 also repositioned towards the acrosome, with chromosomes 8, 9, and Y localized to the periphery. These findings align with previous research suggesting that chromosomal size and gene density influence their own spatial organization within the sperm nucleus. Chromosome 4, being one of the larger human chromosomes, may be more susceptible to mechanical forces during topological reorganization (145–147). It was also observed that chromosome 4 and 8 colocalized and assumed a position in the middle part of sperm nucleus in non-fractionated ejaculate, SU and BPG sperm fractions, as in previously published topological studies (23,146). Chromosomes 7 and 9 have been shown to localize primarily in the middle or peripheral regions of the sperm nucleus (12,146), and their stable positioning in the central nuclear regions also in our study may suggest their role in maintaining nuclear architecture.

We further observed that chromosomes 18, X, and Y align in a consistent order across all the analyzed sperm fractions, from the sperm tail attachment point to the acrosome, following the sequence: 18, Y, and X, as in previous studies (23,145). This order might reflect functional significance, potentially linked to chromosomal properties such as size, gene density, or their involvement in early embryonic development (35). Furthermore, in SU sperm fraction, chromosome 18 in spermatozoa carrying the X chromosome shifts its position, and chromosome X also changes its position, resulting in no change in the distance between these chromosomes. In contrast, in spermatozoa carrying the Y chromosome, chromosome 18 remains in unchanged position, while chromosome Y shifts, leading to an increase in the distance between these chromosomes. Our observation that the positioning of chromosome 18, a gene-poor chromosome, towards the nuclear periphery is consistent with earlier studies indicating a preferential peripheral localization for smaller, gene-poor chromosomes (12,136,145,146). The observed pattern supports the theory that chromosomes with fewer genes may be positioned in regions of the nucleus that are less transcriptionally active after fertilization, reflecting their low contribution to early embryonic gene expression (148).

Our findings also show that sex chromosomes in high-quality spermatozoa tend to take a position in the apical area of the nucleus known as crucial for initial ooplasmic interaction during fertilization (12,27,103,105,106,112,138,140,143,149,150). Our finding is compatible to previous studies with similar apical localization of sex chromosomes, which may contribute to early transcriptional and chromatin reorganization of the paternal genome during the early stages of embryonal development adding some kind of the epigenetic layer (12,27,103,105,106,112,137,138,140,142,143,146,149–155). What is important, previously reported alterations in the positioning of sex chromosomes have been documented in cases with severe types of infertility, in samples with reduced semen quality, such as low sperm count or motility (127). In our study, also the centromere of chromosome Y shifted towards the peripheral region in SU sperm fraction, in contrast to non-fractionated ejaculate. In previous studies, chromosome Y showed variability in centromere positioning, which may reflect the smaller size, lower gene density, and specific gene functions of the Y chromosome (23,105,106,148). Chromosome Y is known to exhibit the most interindividual differences in the localization within sperm cell nucleus (23,105,106,112). In our study, it was shown that both fractioning methods effectively reduced the variability of Y chromosome positioning. What is interesting, chromosome 18 in Y-bearing spermatozoa, showed minimal positional variability, observed across analysed sperm fractions samples, what seems to highlight the independent positioning pattern of chromosomes 18 and Y. Interestingly, in X-bearing sperm, positioning variability of chromosome 18 increased in both good-quality fractions, while variability of chromosome X positioning remained moderate. The rising variability of chromosome 18 in X-bearing sperm cells, alongside consistent X chromosome positioning and unchanged distances between them, may point to a some kind of linkage between reciprocal positioning between these chromosomes within sperm cell nucleus.

In our study an interesting issue was observed for K2 case, in which fractionation improved sperm chromatin protamination status the most, among all evaluated cases. Simultaneously, only in K2 observed chromosome positioning shifted between native ejaculated sample and both good-quality sperm fractions, for each of the evaluated chromosomes (Additional file 6). These observations seem to clearly point to the reasonable application of sperm fractioning methods and their role in enhancing both chromatin quality and chromosomal topology.

The choice of sperm fractioning methods for this study was driven also by their relevance to assisted reproductive technologies (ART), particularly *in vitro* fertilization (IVF) and intracytoplasmic sperm injection (ICSI), where initial fractioning of ejaculate is often applied to improve fertilization outcomes. However, ICSI has raised concerns because it bypasses natural selection mechanisms, potentially increasing the risk of injecting sperm with chromosomal aberrations or fragmented DNA, which are not detectable under standard microscopic evaluation by the embryologist (98,156). One of the major differences between natural fertilization vs. ICSI is the absence of sperm modifications (e.g. acrosomal reaction, membrane fusion), which occur at the sperm entry to the oocyte, which can lead to delayed decondensation of the sub-acrosomal nuclear region in the sperm cell (155). This is associated with increased fertilization abnormalities and failures following ICSI (12,155). Moreover, human sperm injected into hamster eggs also showed that the aberrant nuclear decondensation started from the basal region (155). If sperm nuclei undergo delayed or hindered decondensation during male pronucleus development, the apical localization of the sex chromosomes may contribute to an increased risk of sex chromosome aberrations in ICSI-conceived offspring. This could occur due to delayed entry into the S-phase, which may result in mitotic errors during the first cleavage (28,155). This hypothesis can be supported by the fact that an increased rate of sex chromosomal aberrations has been reported in progeny conceived by ICSI (157,158). It is interesting, that in our study, the X and Y chromosomes in the good-quality sperm fractions did not move toward the acrosome but instead localized to the nuclear periphery, seems to be elevating and fortifying the importance of the chromosomal organization context. This hypothesis requires further investigation, including also other methods of sperm selection, as well as high resolution mapping of the sperm genome and epigenome based on chromatin capture basis.

## Conclusions

In conclusion, our study demonstrated that high-quality sperm selection, based on motility and morphology, significantly enhanced chromatin protamination, reduced sperm ssDNA fragmentation, increased 5mC and 5hmC levels of global sperm DNA, and showed distinct patterns of chromosome positioning. Chromosome repositioning was documented, particularly for sex chromosomes, which shifted towards the sperm nuclear periphery in the apical region, which is crucial for the initial interaction with the ooplasm during fertilization. The findings highlight the importance of sperm fractionation methods in the chromosomal and chromatin context concerning selection of spermatozoa with good characteristics for assisted reproductive technologies (ART). This targeted selection highlights the importance of further research on chromatin interactions and dynamics in spermatozoa, and their implications for fertilization and embryo development, also to refine ART protocols for improved outcomes.

## Supporting information

List of Additional files

## List of abbreviations

5hmC DNA: hydroxymethylation
5mC DNA: methylation
AB: Aniline blue
AO: Acridine orange
ART: Assisted reproductive technology
BGR: Blue, green and red filter
BPG: Bottom layer of Percoll gradient
BSA: Bovine serum albumin
ChIA-Drop: Chromatin interaction analysis with droplet-based high-throughput sequencing
CT: Chromosome territory
DAPI: 4′,6-diamidino-2-phenylindole ds
DNA: double-stranded DNA
DTT: Dithiothreitol
FISH: Fluorescent *in situ* hybridization
GAM: Genome architecture mapping
Hi-C: High-throughput chromosome conformation capture
ICSI: Intracytoplasmic Sperm Injection
IF: Immunofluorescence staining
IMSI: Intracytoplasmic Morphologically Selected Sperm Injection
IVF: *In vitro* Fertilization
PBS: Phosphate-buffered saline
PBST: PBS + Triton X-100
SpG: Spectrum Green
SpO: Spectrum Orange
SPRITE: Split-Pool Recognition of Interactions by Tag Extension
SSC: Saline sodium citrate
ssDNA: single-stranded
DNA SU: Swim up fraction
TAD: Topologically associating domain
Tet Ten-Eleven: Translocation Proteins
TLC: Thin-layer chromatography
TP: transition protein
WHO: World Health Organization
XCI: X chromosome inactivation

## Supplementary Materials

### Declarations

**Funding:** This work was funded by the National Science Centre in Poland, grant no.: 2020/38/E/NZ2/00134 (to MO).

**Institutional Review Board Statement:** Ethical Committee approval (Local Bioethical Committee at Poznan University of Medical Sciences, approval No. 669/22) was received for the study. All participants were notified about the aim of the study, and provided written informed consent. All experiments were performed in accordance with relevant guidelines and regulations.

**Informed Consent Statement:** Informed consent was obtained from all subjects involved in the study.

**Consent for publication:** No personal data of any of the participant in this study are provided. Hence, no identification of the participants is possible. Nevertheless, written informed consent was obtained from all men.

**Data Availability Statement:** All the data generated during this study are included in this manuscript, its Supplementary Information Files and deposited in a Zenodo repository (DOI: 10.5281/zenodo.15599300).

**Author Contributions: ZG** collection of data, samples preparation, evaluation of all results, data interpretation, manuscript drafting; **JK**, **JP**, **ZM** help in data collection and samples preparation; **MKa** semen analysis and selection of samples; **MF** TUNEL assay evaluation; **MO** conceptualization, supervision of the work, data interpretation, revision and finalization of the manuscript, funds collection; **MKu** data interpretation and critical revision of the manuscript.

## Acknowledgments

We would like to thank our M.Sc. students – Martyna Zurek and Aleksandra Zasina, for their partial help in measurements of topology data.

## References

1. WHO laboratory manual for the examination and processing of human semen. 2021. Available from: https://www.who.int/publications-detail-redirect/9789240030787

2. Selvaraju V, Baskaran S, Agarwal A, Henkel R. Environmental contaminants and male infertility: Effects and mechanisms. Andrologia. 2021 Feb;53(1):e13646.

3. Ferlin A, Raicu F, Gatta V, Zuccarello D, Palka G, Foresta C. Male infertility: role of genetic background. Reprod Biomed Online. 2007 Jun;14(6):734–45.

4. Hamada A, Esteves SC, Nizza M, Agarwal A. Unexplained male infertility: diagnosis and management. Int Braz J Urol. 2012;38(5):576–94.

5. Krausz C, Riera-Escamilla A. Genetics of male infertility. Nat Rev Urol. 2018 Jun;15(6):369–84.

6. Colaco S, Sakkas D. Paternal factors contributing to embryo quality. J Assist Reprod Genet. 2018 Nov;35(11):1953–68.

7. Dadoune JP. Spermatozoal RNAs: what about their functions? Microsc Res Tech. 2009 Aug;72(8):536–51.

8. Ioannou D, Millan NM, Jordan E, Tempest HG. A new model of sperm nuclear architecture following assessment of the organization of centromeres and telomeres in three-dimensions. Sci Rep. 2017 Jan 31;7:41585.

9. Oliva R. Protamines and male infertility. Human Reproduction Update. 2006 Aug 1;12(4):417– 35.

10. Zhang X, Gabriel MS, Zini A. Sperm Nuclear Histone to Protamine Ratio in Fertile and Infertile Men: Evidence of Heterogeneous Subpopulations of Spermatozoa in the Ejaculate. Journal of Andrology. 2006;27(3):414–20.

11. Ainsworth C. Cell biology: the secret life of sperm. Nature. 2005 Aug 11;436(7052):770–1.

12. Zalenskaya IA, Zalensky AO. Non-random positioning of chromosomes in human sperm nuclei. Chromosome Res. 2004;12(2):163–73.

13. Hammoud SS, Nix DA, Zhang H, Purwar J, Carrell DT, Cairns BR. Distinctive chromatin in human sperm packages genes for embryo development. Nature. 2009 Jul 23;460(7254):473–8.

14. Ward WS. Function of sperm chromatin structural elements in fertilization and development. Mol Hum Reprod. 2010 Jan;16(1):30–6.

15. Jung YH, Sauria MEG, Lyu X, Cheema MS, Ausio J, Taylor J, et al. Chromatin States in Mouse Sperm Correlate with Embryonic and Adult Regulatory Landscapes. Cell Rep. 2017 Feb 7;18(6):1366–82.

16. Steger K, Wilhelm J, Konrad L, Stalf T, Greb R, Diemer T, et al. Both protamine-1 to protamine-2 mRNA ratio and Bcl2 mRNA content in testicular spermatids and ejaculated spermatozoa discriminate between fertile and infertile men. Human Reproduction. 2008 Jan 1;23(1):11–6.

17. Carrell DT, Liu L. Altered protamine 2 expression is uncommon in donors of known fertility, but common among men with poor fertilizing capacity, and may reflect other abnormalities of spermiogenesis. J Androl. 2001;22(4):604–10.

18. Nasr-Esfahani MH, Razavi S, Mozdarani H, Mardani M, Azvagi H. Relationship between protamine deficiency with fertilization rate and incidence of sperm premature chromosomal condensation post-ICSI. Andrologia. 2004;36(3):95–100.

19. Tena JJ, Santos-Pereira JM. Topologically Associating Domains and Regulatory Landscapes in Development, Evolution and Disease. Front Cell Dev Biol. 2021;9:702787.

20. Lupiáñez DG, Kraft K, Heinrich V, Krawitz P, Brancati F, Klopocki E, et al. Disruptions of topological chromatin domains cause pathogenic rewiring of gene-enhancer interactions. Cell. 2015 May 21;161(5):1012–25.

21. Spielmann M, Lupiáñez DG, Mundlos S. Structural variation in the 3D genome. Nat Rev Genet. 2018 Jul;19(7):453–67.

22. Kempfer R, Pombo A. Methods for mapping 3D chromosome architecture. Nat Rev Genet. 2020 Apr;21(4):207–26.

23. Olszewska M, Wiland E, Huleyuk N, Fraczek M, Midro AT, Zastavna D, et al. Chromosome (re)positioning in spermatozoa of fathers and sons - carriers of reciprocal chromosome translocation (RCT). BMC Med Genomics. 2019 Feb 1;12(1):30.

24. Schuck PL, Stewart JA. FISHing for Damage on Metaphase Chromosomes. Methods Mol Biol. 2019;1999:335–47.

25. Zalensky AO, Allen MJ, Kobayashi A, Zalenskaya IA, Balhórn R, Bradbury EM. Well-defined genome architecture in the human sperm nucleus. Chromosoma. 1995 May;103(9):577–90.

26. Wiland E, Fraczek M, Olszewska M, Kurpisz M. Topology of chromosome centromeres in human sperm nuclei with high levels of DNA damage. Sci Rep. 2016 Aug 25;6:31614.

27. Hazzouri M, Rousseaux S, Mongelard F, Usson Y, Pelletier R, Faure AK, et al. Genome organization in the human sperm nucleus studied by FISH and confocal microscopy. Mol Reprod Dev. 2000 Mar;55(3):307–15.

28. Luetjens CM, Payne C, Schatten G. Non-random chromosome positioning in human sperm and sex chromosome anomalies following intracytoplasmic sperm injection. Lancet. 1999 Apr 10;353(9160):1240.

29. Jones EL, Mudrak O, Zalensky AO. Kinetics of human male pronuclear development in a heterologous ICSI model. J Assist Reprod Genet. 2010 Jun;27(6):277–83.

30. Lowe R, Gemma C, Rakyan VK, Holland ML. Sexually dimorphic gene expression emerges with embryonic genome activation and is dynamic throughout development. BMC Genomics. 2015 Apr 14;16(1):295.

31. Werner RJ, Schultz BM, Huhn JM, Jelinek J, Madzo J, Engel N. Sex chromosomes drive gene expression and regulatory dimorphisms in mouse embryonic stem cells. Biol Sex Differ. 2017 Aug 17;8(1):28.

32. Schulz EG, Meisig J, Nakamura T, Okamoto I, Sieber A, Picard C, et al. The two active X chromosomes in female ESCs block exit from the pluripotent state by modulating the ESC signaling network. Cell Stem Cell. 2014 Feb 6;14(2):203–16.

33. Chen G, Schell JP, Benitez JA, Petropoulos S, Yilmaz M, Reinius B, et al. Single-cell analyses of X Chromosome inactivation dynamics and pluripotency during differentiation. Genome Res. 2016 Oct;26(10):1342–54.

34. Zhou Q, Wang T, Leng L, Zheng W, Huang J, Fang F, et al. Single-cell RNA-seq reveals distinct dynamic behavior of sex chromosomes during early human embryogenesis. Mol Reprod Dev. 2019 Jul;86(7):871–82.

35. Richardson V, Engel N, Kulathinal RJ. Comparative developmental genomics of sex-biased gene expression in early embryogenesis across mammals. Biol Sex Differ. 2023 May 19;14(1):30.

36. Engel N. Sex Differences in Early Embryogenesis: Inter-Chromosomal Regulation Sets the Stage for Sex-Biased Gene Networks: The dialogue between the sex chromosomes and autosomes imposes sexual identity soon after fertilization. Bioessays. 2018 Sep;40(9):e1800073.

37. Blencowe M, Chen X, Zhao Y, Itoh Y, McQuillen CN, Han Y, et al. Relative contributions of sex hormones, sex chromosomes, and gonads to sex differences in tissue gene regulation. Genome Res. 2022 May;32(5):807–24.

38. Barakat TS, Gribnau J. X chromosome inactivation in the cycle of life. Development. 2012 Jun;139(12):2085–9.

39. Latham KE. X chromosome imprinting and inactivation in preimplantation mammalian embryos. Trends Genet. 2005 Feb;21(2):120–7.

40. Lee JT. Functional intergenic transcription: a case study of the X-inactivation centre. Philos Trans R Soc Lond B Biol Sci. 2003 Aug 29;358(1436):1417–23; discussion 1423.

41. Migeon BR. X chromosome inactivation: theme and variations. Cytogenet Genome Res. 2002;99(1–4):8–16.

42. Plath K, Mlynarczyk-Evans S, Nusinow DA, Panning B. Xist RNA and the mechanism of X chromosome inactivation. Annu Rev Genet. 2002;36:233–78.

43. Disteche CM, Berletch JB. X-chromosome inactivation and escape. J Genet. 2015 Dec;94(4):591–9.

44. van den Berg IM, Laven JSE, Stevens M, Jonkers I, Galjaard RJ, Gribnau J, et al. X chromosome inactivation is initiated in human preimplantation embryos. Am J Hum Genet. 2009 Jun;84(6):771–9.

45. Tan K, An L, Miao K, Ren L, Hou Z, Tao L, et al. Impaired imprinted X chromosome inactivation is responsible for the skewed sex ratio following in vitro fertilization. Proceedings of the National Academy of Sciences. 2016 Mar 22;113(12):3197–202.

46. Lanasa MC, Hogge WA, Kubik C, Blancato J, Hoffman EP. Highly Skewed X-Chromosome Inactivation Is Associated with Idiopathic Recurrent Spontaneous Abortion. The American Journal of Human Genetics. 1999 Jul 1;65(1):252–4.

47. Sullivan AE, Lewis T, Stephenson M, Odem R, Schreiber J, Ober C, et al. Pregnancy outcome in recurrent miscarriage patients with skewed X chromosome inactivation. Obstet Gynecol. 2003 Jun;101(6):1236–42.

48. Petropoulos S, Edsgärd D, Reinius B, Deng Q, Panula SP, Codeluppi S, et al. Single-Cell RNA-Seq Reveals Lineage and X Chromosome Dynamics in Human Preimplantation Embryos. Cell. 2016 Sep 22;167(1):285.

49. Fisher EMC, Beer-Romero P, Brown LG, Ridley A, McNeil JA, Lawrence JB, et al. Homologous ribosomal protein genes on the human X and Y chromosomes: Escape from X inactivation and possible implications for turner syndrome. Cell. 1990 Dec 21;63(6):1205–18.

50. Andrés O, Kellermann T, López-Giráldez F, Rozas J, Domingo-Roura X, Bosch M. RPS4Ygene family evolution in primates. BMC Evolutionary Biology. 2008 May 13;8(1):142.

51. Musio A. The multiple facets of the SMC1A gene. Gene. 2020 Jun 15;743:144612.

52. Yatskevich S, Rhodes J, Nasmyth K. Organization of Chromosomal DNA by SMC Complexes. Annu Rev Genet. 2019 Dec 3;53:445–82.

53. Jessberger R, Podust V, Hübscher U, Berg P. A mammalian protein complex that repairs double-strand breaks and deletions by recombination. J Biol Chem. 1993 Jul 15;268(20):15070–9.

54. Jessberger R, Riwar B, Baechtold H, Akhmedov AT. SMC proteins constitute two subunits of the mammalian recombination complex RC-1. EMBO J. 1996 Aug 1;15(15):4061–8.

55. Musio A, Montagna C, Zambroni D, Indino E, Barbieri O, Citti L, et al. Inhibition of BUB1 results in genomic instability and anchorage-independent growth of normal human fibroblasts. Cancer Res. 2003 Jun 1;63(11):2855–63.

56. Musio A, Montagna C, Mariani T, Tilenni M, Focarelli ML, Brait L, et al. SMC1 involvement in fragile site expression. Hum Mol Genet. 2005 Feb 15;14(4):525–33.

57. Xiang JF, Corces VG. Regulation of 3D chromatin organization by CTCF. Curr Opin Genet Dev. 2021 Apr;67:33–40.

58. Lismer A, Kimmins S. Emerging evidence that the mammalian sperm epigenome serves as a template for embryo development. Nat Commun. 2023 Apr 14;14(1):2142.

59. Rao SSP, Huntley MH, Durand NC, Stamenova EK, Bochkov ID, Robinson JT, et al. A 3D map of the human genome at kilobase resolution reveals principles of chromatin looping. Cell. 2014 Dec 18;159(7):1665–80.

60. Schwarzer W, Abdennur N, Goloborodko A, Pekowska A, Fudenberg G, Loe-Mie Y, et al. Two independent modes of chromatin organization revealed by cohesin removal. Nature. 2017 Nov 2;551(7678):51–6.

61. Butler BA, Soong J, Gergen JP. The Drosophila segmentation gene runt has an extended cis-regulatory region that is required for vital expression at other stages of development. Mech Dev. 1992 Nov;39(1–2):17–28.

62. Sassone-Corsi P. Unique chromatin remodeling and transcriptional regulation in spermatogenesis. Science. 2002 Jun 21;296(5576):2176–8.

63. Wykes SM, Krawetz SA. The structural organization of sperm chromatin. J Biol Chem. 2003 Aug 8;278(32):29471–7.

64. Pepin AS, Jazwiec PA, Dumeaux V, Sloboda DM, Kimmins S. Determining the effects of paternal obesity on sperm chromatin at histone H3 lysine 4 tri-methylation in relation to the placental transcriptome and cellular composition. Elife. 2024 Nov 29;13:e83288.

65. Odroniec A, Olszewska M, Kurpisz M. Epigenetic markers in the embryonal germ cell development and spermatogenesis. Basic and Clinical Andrology. 2023 Feb 23;33(1):6.

66. Sujit KM, Sarkar S, Singh V, Pandey R, Agrawal NK, Trivedi S, et al. Genome-wide differential methylation analyses identifies methylation signatures of male infertility. Hum Reprod. 2018 Dec 1;33(12):2256–67.

67. Xavier MJ, Roman SD, Aitken RJ, Nixon B. Transgenerational inheritance: how impacts to the epigenetic and genetic information of parents affect offspring health. Hum Reprod Update. 2019 Sep 11;25(5):518–40.

68. Castillo J, Jodar M, Oliva R. The contribution of human sperm proteins to the development and epigenome of the preimplantation embryo. Human Reproduction Update. 2018 Sep 1;24(5):535–55.

69. Yu B, Zhou H, Liu M, Zheng T, Jiang L, Zhao M, et al. Epigenetic Alterations in Density Selected Human Spermatozoa for Assisted Reproduction. PLOS ONE. 2015 Dec 28;10(12):e0145585.

70. Yan R, Cheng X, Gu C, Xu Y, Long X, Zhai J, et al. Dynamics of DNA hydroxymethylation and methylation during mouse embryonic and germline development. Nat Genet. 2023 Jan;55(1):130–43.

71. Stewart KR, Veselovska L, Kelsey G. Establishment and Functions of DNA Methylation in the Germline. Epigenomics. 2016 Oct 1;8(10):1399–413.

72. Ge SQ, Lin SL, Zhao ZH, Sun QY. Epigenetic dynamics and interplay during spermatogenesis and embryogenesis: implications for male fertility and offspring health. Oncotarget. 2017 Aug 8;8(32):53804–18.

73. Monk D, Mackay DJG, Eggermann T, Maher ER, Riccio A. Genomic imprinting disorders: lessons on how genome, epigenome and environment interact. Nat Rev Genet. 2019 Apr;20(4):235–48.

74. Puri D, Dhawan J, Mishra RK. The paternal hidden agenda: Epigenetic inheritance through sperm chromatin. Epigenetics. 2010 Jul 1;5(5):386–91.

75. Lambrot R, Xu C, Saint-Phar S, Chountalos G, Cohen T, Paquet M, et al. Low paternal dietary folate alters the mouse sperm epigenome and is associated with negative pregnancy outcomes. Nat Commun. 2013 Dec 10;4(1):2889.

76. Kaati G, Bygren LO, Edvinsson S. Cardiovascular and diabetes mortality determined by nutrition during parents’ and grandparents’ slow growth period. Eur J Hum Genet. 2002 Nov;10(11):682– 8.

77. Wei Y, Yang CR, Wei YP, Zhao ZA, Hou Y, Schatten H, et al. Paternally induced transgenerational inheritance of susceptibility to diabetes in mammals. Proceedings of the National Academy of Sciences. 2014 Feb 4;111(5):1873–8.

78. Liang F, Diao L, Liu J, Jiang N, Zhang J, Wang H, et al. Paternal ethanol exposure and behavioral abnormities in offspring: Associated alterations in imprinted gene methylation. Neuropharmacology. 2014 Jun 1;81:126–33.

79. Dias BG, Ressler KJ. Parental olfactory experience influences behavior and neural structure in subsequent generations. Nat Neurosci. 2014 Jan;17(1):89–96.

80. Meyer RG, Ketchum CC, Meyer-Ficca ML. Heritable sperm chromatin epigenetics: a break to remember. Biol Reprod. 2017 Jan 1;97(6):784–97.

81. Allis CD, Jenuwein T. The molecular hallmarks of epigenetic control. Nat Rev Genet. 2016 Aug;17(8):487–500.

82. Luo C, Hajkova P, Ecker JR. Dynamic DNA methylation: In the right place at the right time. Science. 2018 Sep 28;361(6409):1336–40.

83. Allegrucci C, Thurston A, Lucas E, Young L. Epigenetics and the germline. Reprodaction. 2005 Feb 1;129(2):137–149.

84. Mayer W, Niveleau A, Walter J, Fundele R, Haaf T. Demethylation of the zygotic paternal genome. Nature. 2000 Feb 3;403(6769):501–2.

85. Aston KI, Punj V, Liu L, Carrell DT. Genome-wide sperm deoxyribonucleic acid methylation is altered in some men with abnormal chromatin packaging or poor in vitro fertilization embryogenesis. Fertility and Sterility. 2012 Feb 1;97(2):285–292.e4.

86. Carrell DT. Epigenetics of the male gamete. Fertility and Sterility. 2012 Feb 1;97(2):267–74.

87. Ito S, D’Alessio AC, Taranova OV, Hong K, Sowers LC, Zhang Y. Role of Tet proteins in 5mC to 5hmC conversion, ES-cell self-renewal and inner cell mass specification. Nature. 2010 Aug 26;466(7310):1129–33.

88. Delatte B, Fuks F. TET proteins: on the frenetic hunt for new cytosine modifications. Brief Funct Genomics. 2013 May;12(3):191–204.

89. Ecsedi S, Rodríguez-Aguilera JR, Hernandez-Vargas H. 5-Hydroxymethylcytosine (5hmC), or How to Identify Your Favorite Cell. Epigenomes. 2018 Mar;2(1):3.

90. Xu T, Gao H. Hydroxymethylation and tumors: can 5-hydroxymethylation be used as a marker for tumor diagnosis and treatment? Human Genomics. 2020 May 6;14(1):15.

91. Guz J, Gackowski D, Foksinski M, Rozalski R, Olinski R. Comparison of the Absolute Level of Epigenetic Marks 5-Methylcytosine, 5-Hydroxymethylcytosine, and 5-Hydroxymethyluracil Between Human Leukocytes and Sperm1. Biology of Reproduction. 2014 Sep 1;91(3):55, 1–5.

92. Tahiliani M, Koh KP, Shen Y, Pastor WA, Bandukwala H, Brudno Y, et al. Conversion of 5- Methylcytosine to 5-Hydroxymethylcytosine in Mammalian DNA by MLL Partner TET1. Science. 2009 May 15;324(5929):930–5.

93. Nabel CS, Jia H, Ye Y, Shen L, Goldschmidt HL, Stivers JT, et al. AID/APOBEC deaminases disfavor modified cytosines implicated in DNA demethylation. Nat Chem Biol. 2012 Sep;8(9):751–8.

94. Szyf M. The Elusive Role of 5′-Hydroxymethylcytosine. Epigenomics. 2016 Nov 1;8(11):1539–51.

95. Rao M, Tang L, Wang L, Chen M, Yan G, Zhao S. Cumulative live birth rates after IVF/ICSI cycles with sperm prepared by density gradient centrifugation vs. swim-up: a retrospective study using a propensity score-matching analysis. Reprod Biol Endocrinol. 2022 Mar 31;20(1):60.

96. Baldini D, Ferri D, Baldini GM, Lot D, Catino A, Vizziello D, et al. Sperm Selection for ICSI: Do We Have a Winner? Cells. 2021 Dec;10(12):3566.

97. Cassuto NG, Ogal N, Assou S, Ruoso L, Rogers EJ, Monteiro MJ, et al. Different Nuclear Architecture in Human Sperm According to Their Morphology. Genes. 2024 Apr 7;15(4):464.

98. Nasr-Esfahani MH, Salehi M, Razavi S, Anjomshoa M, Rozbahani S, Moulavi F, et al. Effect of sperm DNA damage and sperm protamine deficiency on fertilization and embryo development post-ICSI. Reprod Biomed Online. 2005 Aug;11(2):198–205.

99. Canale D, Giorgi PM, Gasperini M, Pucci E, Barletta D, Gasperi M, et al. Inter and intra-individual variability of sperm morphology after selection with three different techniques: layering, swimup from pellet and percoll. J Endocrinol Invest. 1994 Oct;17(9):729–32.

100. Olszewska M, Huleyuk N, Fraczek M, Zastavna D, Wiland E, Kurpisz M. Sperm FISH and chromatin integrity in spermatozoa from a t(6;10;11) carrier. Reproduction. 2014 May;147(5):659–70.

101. Olszewska M, Kordyl O, Kamieniczna M, Fraczek M, Jędrzejczak P, Kurpisz M. Global 5mC and 5hmC DNA Levels in Human Sperm Subpopulations with Differentially Protaminated Chromatin in Normo- and Oligoasthenozoospermic Males. Int J Mol Sci. 2022 Apr 19;23(9):4516.

102. Olszewska M, Barciszewska MZ, Fraczek M, Huleyuk N, Chernykh VB, Zastavna D, et al. Global methylation status of sperm DNA in carriers of chromosome structural aberrations. Asian Journal of Andrology. 2017 Feb;19(1):117.

103. Olszewska M, Wanowska E, Kishore A, Huleyuk N, Georgiadis AP, Yatsenko AN, et al. Genetic dosage and position effect of small supernumerary marker chromosome (sSMC) in human sperm nuclei in infertile male patient. Sci Rep. 2015 Nov 30;5:17408.

104. Olszewska M, Fraczek M, Huleyuk N, Czernikiewicz A, Wiland E, Boksa M, et al. Chromatin structure analysis of spermatozoa from reciprocal chromosome translocation (RCT) carriers with known meiotic segregation patterns. Reprod Biol. 2013 Sep;13(3):209–20.

105. Olszewska M, Wiland E, Kurpisz M. Positioning of chromosome 15, 18, X and Y centromeres in sperm cells of fertile individuals and infertile patients with increased level of aneuploidy. Chromosome Res. 2008;16(6):875–90.

106. Wiland E, Zegało M, Kurpisz M. Interindividual differences and alterations in the topology of chromosomes in human sperm nuclei of fertile donors and carriers of reciprocal translocations. Chromosome Res. 2008;16(2):291–305.

107. Wiland E, Olszewska M, Georgiadis A, Huleyuk N, Panasiuk B, Zastavna D, et al. Cytogenetic and molecular analyses of de novo translocation dic(9;13)(p11.2;p12) in an infertile male. Mol Cytogenet. 2014 Feb 21;7(1):14.

108. Wiland E, Yatsenko AN, Kishore A, Stanczak H, Zdarta A, Ligaj M, et al. FISH and array CGH characterization of de novo derivative Y chromosome (Yq duplication and partial Yp deletion) in an azoospermic male. Reprod Biomed Online. 2015 Aug;31(2):217–24.

109. Wiland E, Olszewska M, Huleyuk N, Chernykh VB, Kurpisz M. The effect of Robertsonian translocations on the intranuclear positioning of NORs (nucleolar organizing regions) in human sperm cells. Sci Rep. 2019 Feb 18;9(1):2213.

110. Olszewska M, Stokowy T, Pollock N, Huleyuk N, Georgiadis A, Yatsenko S, et al. Familial Infertility (Azoospermia and Cryptozoospermia) in Two Brothers—Carriers of t(1;7) Complex Chromosomal Rearrangement (CCR): Molecular Cytogenetic Analysis. International Journal of Molecular Sciences. 2020 Jan;21(12):4559.

111. Efimova OA, Pendina AA, Tikhonov AV, Parfenyev SE, Mekina ID, Komarova EM, et al. Genome-wide 5-hydroxymethylcytosine patterns in human spermatogenesis are associated with semen quality. Oncotarget. 2017 Jun 1;8(51):88294–307.

112. Wiland E, Fraczek M, Olszewska M, Kurpisz M. Topology of chromosome centromeres in human sperm nuclei with high levels of DNA damage. Sci Rep. 2016 Aug 25;6:31614.

113. Ribas-Maynou J, Novo S, Salas-Huetos A, Rovira S, Antich M, Yeste M. Condensation and protamination of sperm chromatin affect ICSI outcomes when gametes from healthy individuals are used. Hum Reprod. 2023 Mar 1;38(3):371–86.

114. Yoshida K, Muratani M, Araki H, Miura F, Suzuki T, Dohmae N, et al. Mapping of histone-binding sites in histone replacement-completed spermatozoa. Nat Commun. 2018 Sep 24;9(1):3885.

115. Hammoud SS, Purwar J, Pflueger C, Cairns BR, Carrell DT. Alterations in sperm DNA methylation patterns at imprinted loci in two classes of infertility. Fertility and Sterility. 2010 Oct 1;94(5):1728–33.

116. Nanassy L, Carrell DT. Analysis of the methylation pattern of six gene promoters in sperm of men with abnormal protamination. Asian J Androl. 2011 Mar;13(2):342–6.

117. Hamad MF. Quantification of histones and protamines mRNA transcripts in sperms of infertile couples and their impact on sperm’s quality and chromatin integrity. Reproductive Biology. 2019 Mar 1;19(1):6–13.

118. Manicardi GC, Bianchi PG, Pantano S, Azzoni P, Bizzaro D, Bianchi U, et al. Presence of endogenous nicks in DNA of ejaculated human spermatozoa and its relationship to chromomycin A3 accessibility. Biol Reprod. 1995 Apr;52(4):864–7.

119. Rajabi H, Mohseni-Kouchesfehani H, Mohammadi-Sangcheshmeh A, Farifteh-Nobijari F, Salehi M. Pronuclear epigenetic modification of protamine deficient human sperm following injection into mouse oocytes. Systems Biology in Reproductive Medicine. 2016 Mar 3;62(2):125–32.

120. Jackson RE, Bormann CL, Hassun PA, Rocha AM, Motta ELA, Serafini PC, et al. Effects of semen storage and separation techniques on sperm DNA fragmentation. Fertility and Sterility. 2010 Dec 1;94(7):2626–30.

121. Donnelly ET, O’Connell M, McClure N, Lewis SE. Differences in nuclear DNA fragmentation and mitochondrial integrity of semen and prepared human spermatozoa. Hum Reprod. 2000 Jul;15(7):1552–61.

122. Marchetti C, Obert G, Deffosez A, Formstecher P, Marchetti P. Study of mitochondrial membrane potential, reactive oxygen species, DNA fragmentation and cell viability by flow cytometry in human sperm. Hum Reprod. 2002 May;17(5):1257–65.

123. Sakkas D, Manicardi GC, Tomlinson M, Mandrioli M, Bizzaro D, Bianchi PG, et al. The use of two density gradient centrifugation techniques and the swim-up method to separate spermatozoa with chromatin and nuclear DNA anomalies. Hum Reprod. 2000 May;15(5):1112–6.

124. Zini A, Finelli A, Phang D, Jarvi K. Influence of semen processing technique on human sperm DNA integrity. Urology. 2000 Dec 20;56(6):1081–4.

125. Gorczyca W, Gong J, Darzynkiewicz Z. Detection of DNA strand breaks in individual apoptotic cells by the in situ terminal deoxynucleotidyl transferase and nick translation assays. Cancer Res. 1993 Apr 15;53(8):1945–51.

126. Evenson D, Jost L. Sperm chromatin structure assay is useful for fertility assessment. Methods Cell Sci. 2000 Jun 1;22(2):169–89.

127. Ioannou D, Meershoek EJ, Christopikou D, Ellis M, Thornhill AR, Griffin DK. Nuclear organisation of sperm remains remarkably unaffected in the presence of defective spermatogenesis. Chromosome Res. 2011 Aug;19(6):741–53.

128. Bazrgar M, Gourabi H, Yazdi PE, Vazirinasab H, Fakhri M, Hassani F, et al. DNA repair signalling pathway genes are overexpressed in poor-quality pre-implantation human embryos with complex aneuploidy. Eur J Obstet Gynecol Reprod Biol. 2014 Apr;175:152–6.

129. Nanassy L, Carrell DT. Abnormal methylation of the promoter of CREM is broadly associated with male factor infertility and poor sperm quality but is improved in sperm selected by density gradient centrifugation. Fertil Steril. 2011 Jun;95(7):2310–4.

130. Laurentino S, Beygo J, Nordhoff V, Kliesch S, Wistuba J, Borgmann J, et al. Epigenetic germline mosaicism in infertile men. Hum Mol Genet. 2015 Mar 1;24(5):1295–304.

131. Montjean D, Zini A, Ravel C, Belloc S, Dalleac A, Copin H, et al. Sperm global DNA methylation level: association with semen parameters and genome integrity. Andrology. 2015 Mar;3(2):235–40.

132. Aston KI, Uren PJ, Jenkins TG, Horsager A, Cairns BR, Smith AD, et al. Aberrant sperm DNA methylation predicts male fertility status and embryo quality. Fertility and Sterility. 2015 Dec 1;104(6):1388–1397.e5.

133. Urdinguio RG, Bayón GF, Dmitrijeva M, Toraño EG, Bravo C, Fraga MF, et al. Aberrant DNA methylation patterns of spermatozoa in men with unexplained infertility. Human Reproduction. 2015 May 1;30(5):1014–28.

134. Benchaib M, Braun V, Ressnikof D, Lornage J, Durand P, Niveleau A, et al. Influence of global sperm DNA methylation on IVF results. Human Reproduction. 2005 Mar 1;20(3):768–73.

135. Benchaïb M, Ajina M, Braun V, Niveleau A, Guérin JF. Méthylation du spermatozoïde en Assistance médicale à la procréation (AMP). Gynécologie Obstétrique & Fertilité. 2006 Sep 1;34(9):836–9.

136. Haaf T, Ward DC. Higher order nuclear structure in mammalian sperm revealed by in situ hybridization and extended chromatin fibers. Exp Cell Res. 1995 Aug;219(2):604–11.

137. Foster HA, Abeydeera LR, Griffin DK, Bridger JM. Non-random chromosome positioning in mammalian sperm nuclei, with migration of the sex chromosomes during late spermatogenesis. J Cell Sci. 2005 May 1;118(Pt 9):1811–20.

138. Perdrix A, Travers A, Clatot F, Sibert L, Mitchell V, Jumeau F, et al. Modification of chromosomal architecture in human spermatozoa with large vacuoles. Andrology. 2013 Jan;1(1):57–66.

139. Finch KA, Fonseka G, Ioannou D, Hickson N, Barclay Z, Chatzimeletiou K, et al. Nuclear organisation in totipotent human nuclei and its relationship to chromosomal abnormality. J Cell Sci. 2008 Mar 1;121(Pt 5):655–63.

140. Finch KA, Fonseka KGL, Abogrein A, Ioannou D, Handyside AH, Thornhill AR, et al. Nuclear organization in human sperm: preliminary evidence for altered sex chromosome centromere position in infertile males. Hum Reprod. 2008 Jun;23(6):1263–70.

141. Gurevitch M, Amiel A, Ben-Zion M, Fejgin M, Bartoov B. Acrocentric centromere organization within the chromocenter of the human sperm nucleus. Mol Reprod Dev. 2001 Dec;60(4):507– 16.

142. Zalensky A, Zalenskaya I. Organization of chromosomes in spermatozoa: an additional layer of epigenetic information? Biochemical Society Transactions. 2007 May 22;35(3):609–11.

143. Alladin N, Moskovtsev SI, Russell H, Kenigsberg S, Lulat AGM, Librach CL. The three-dimensional image analysis of the chromocenter in motile and immotile human sperm. Syst Biol Reprod Med. 2013 Jun;59(3):146–52.

144. Wiland E, Midro AT, Panasiuk B, Kurpisz M. The Analysis of Meiotic Segregation Patterns and Aneuploidy in the Spermatozoa of Father and Son With Translocation t(4;5)(p15.1;p12) and the Prediction of the Individual Probability Rate for Unbalanced Progeny at Birth. Journal of Andrology. 2007;28(2):262–72.

145. Mudrak OS, Nazarov IB, Jones EL, Zalensky AO. Positioning of chromosomes in human spermatozoa is determined by ordered centromere arrangement. PLoS One. 2012;7(12):e52944.

146. Manvelyan M, Hunstig F, Bhatt S, Mrasek K, Pellestor F, Weise A, et al. Chromosome distribution in human sperm - a 3D multicolor banding-study. Mol Cytogenet. 2008 Nov 14;1:25.

147. Tilgen N, Guttenbach M, Schmid M. Heterochromatin is not an adequate explanation for close proximity of interphase chromosomes 1--Y, 9--Y, and 16--Y in human spermatozoa. Exp Cell Res. 2001 May 1;265(2):283–7.

148. Solé M, Blanco J, Gil D, Valero O, Pascual Á, Cárdenas B, et al. Chromosomal positioning in spermatogenic cells is influenced by chromosomal factors associated with gene activity, bouquet formation and meiotic sex chromosome inactivation. Chromosoma. 2021 Sep;130(2– 3):163–75.

149. Karamysheva T, Kosyakova N, Guediche N, Liehr T. Small supernumerary marker chromosomes and the nuclear architecture of sperm - a study in a fertile and an infertile brother. Syst Biol Reprod Med. 2015 Jan;61(1):32–6.

150. Sbracia M, Baldi M, Cao D, Sandrelli A, Chiandetti A, Poverini R, et al. Preferential location of sex chromosomes, their aneuploidy in human sperm, and their role in determining sex chromosome aneuploidy in embryos after ICSI. Hum Reprod. 2002 Feb;17(2):320–4.

151. Brockdorff N. Chromosome silencing mechanisms in X-chromosome inactivation: unknown unknowns. Development. 2011 Dec 1;138(23):5057–65.

152. Greaves IK, Rens W, Ferguson-Smith MA, Griffin D, Marshall Graves JA. Conservation of chromosome arrangement and position of the X in mammalian sperm suggests functional significance. Chromosome Res. 2003 Jul 1;11(5):503–12.

153. McLay DW, Clarke HJ. Remodelling the paternal chromatin at fertilization in mammals. Reproduction. 2003 May;125(5):625–33.

154. Ajduk A, Yamauchi Y, Ward MA. Sperm chromatin remodeling after intracytoplasmic sperm injection differs from that of in vitro fertilization. Biol Reprod. 2006 Sep;75(3):442–51.

155. Terada Y, Luetjens CM, Sutovsky P, Schatten G. Atypical decondensation of the sperm nucleus, delayed replication of the male genome, and sex chromosome positioning following intracytoplasmic human sperm injection (ICSI) into golden hamster eggs: does ICSI itself introduce chromosomal anomalies? Fertil Steril. 2000 Sep;74(3):454–60.

156. Martin RH. The risk of chromosomal abnormalities following ICSI. Hum Reprod. 1996 May;11(5):924–5.

157. Bonduelle M, Liebaers I, Deketelaere V, Derde MP, Camus M, Devroey P, et al. Neonatal data on a cohort of 2889 infants born after ICSI (1991-1999) and of 2995 infants born after IVF (1983-1999). Hum Reprod. 2002 Mar;17(3):671–94.

158. In’t Veld P, Brandenburg H, Verhoeff A, Dhont M, Los F. Sex chromosomal abnormalities and intracytoplasmic sperm injection. Lancet. 1995 Sep 16;346(8977):773.

