## Additional file 1 for "Impact of Sperm Fractionation on Chromosome Positioning, Chromatin Integrity, DNA Methylation and Hydroxymethylation Level"

| Chromosome | Catalogue Number | Locus | Chromosome region | DNA class | Spectrum |
|---|---|---|---|---|---|
| 4 | LPE 004G | D4Z1 | 4p11.1-q11.1 | α-satellite | Green |
| 7 | LPE 007G | D7Z1 | 7p11.1-q11.1 | α-satellite | Green |
| 8 | LPE 008R | D8Z2 | 8p11.1-q11.1 | α-satellite | Orange |
| 9 | LPE 009R | D9Z3 | 9q12 | satellite III | Orange |
| 18 | LPA 004 | D18Z1 | 18p11.1-q11.1 | α-satellite | Aqua |
| X | LPE 0XG | DXZ1 | Xp11.1-q11.1 | α-satellite | Green |
| Y | LPE 0YcR | DYZ3 | Yp11.1-q11.1 | α-satellite | Orange |
