## Supplementary figures and images for "Impact of Sperm Fractionation on Chromosome Positioning, Chromatin Integrity, DNA Methylation and Hydroxymethylation Level"

### Additional file 5

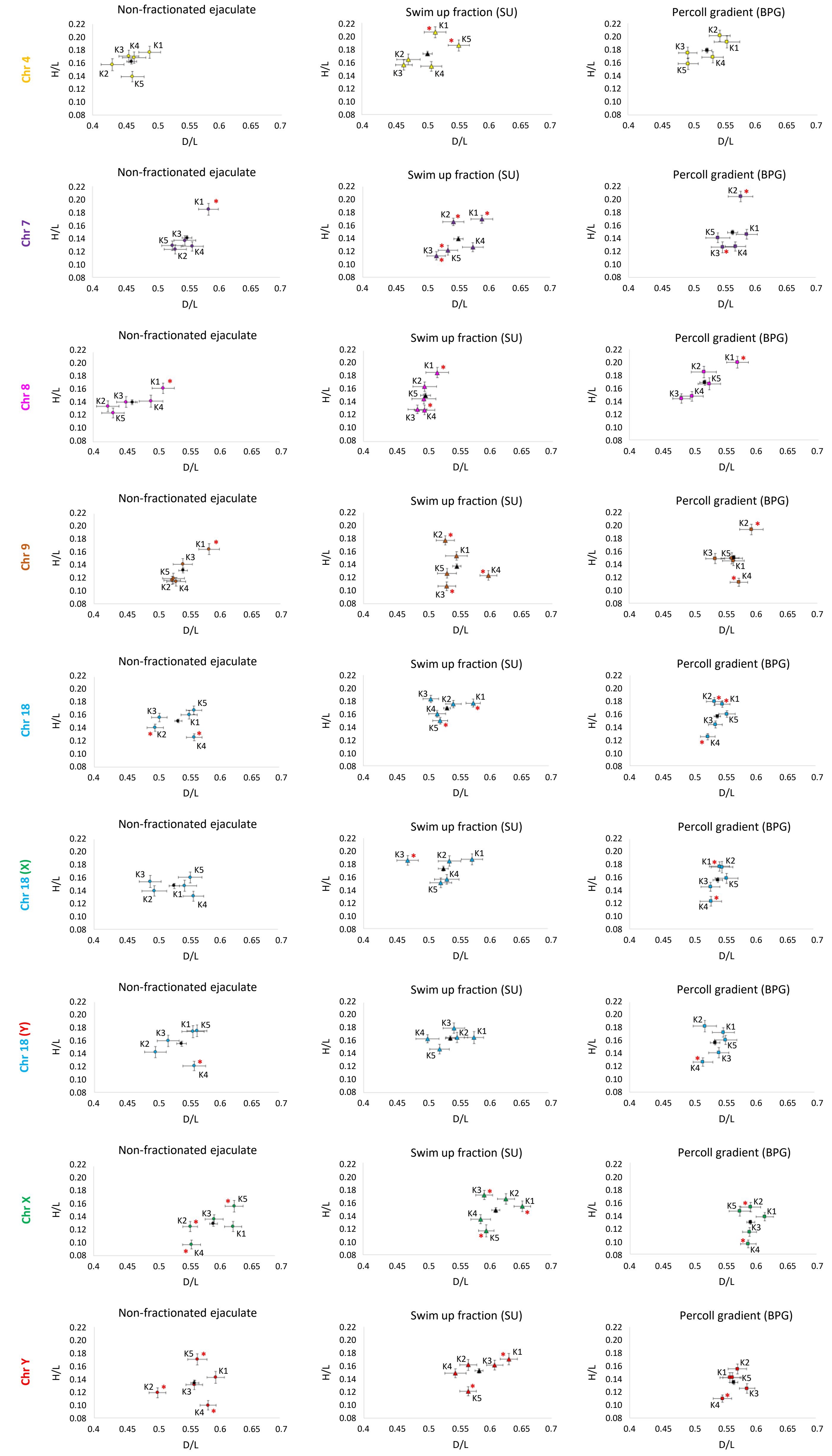

### Additional file 6

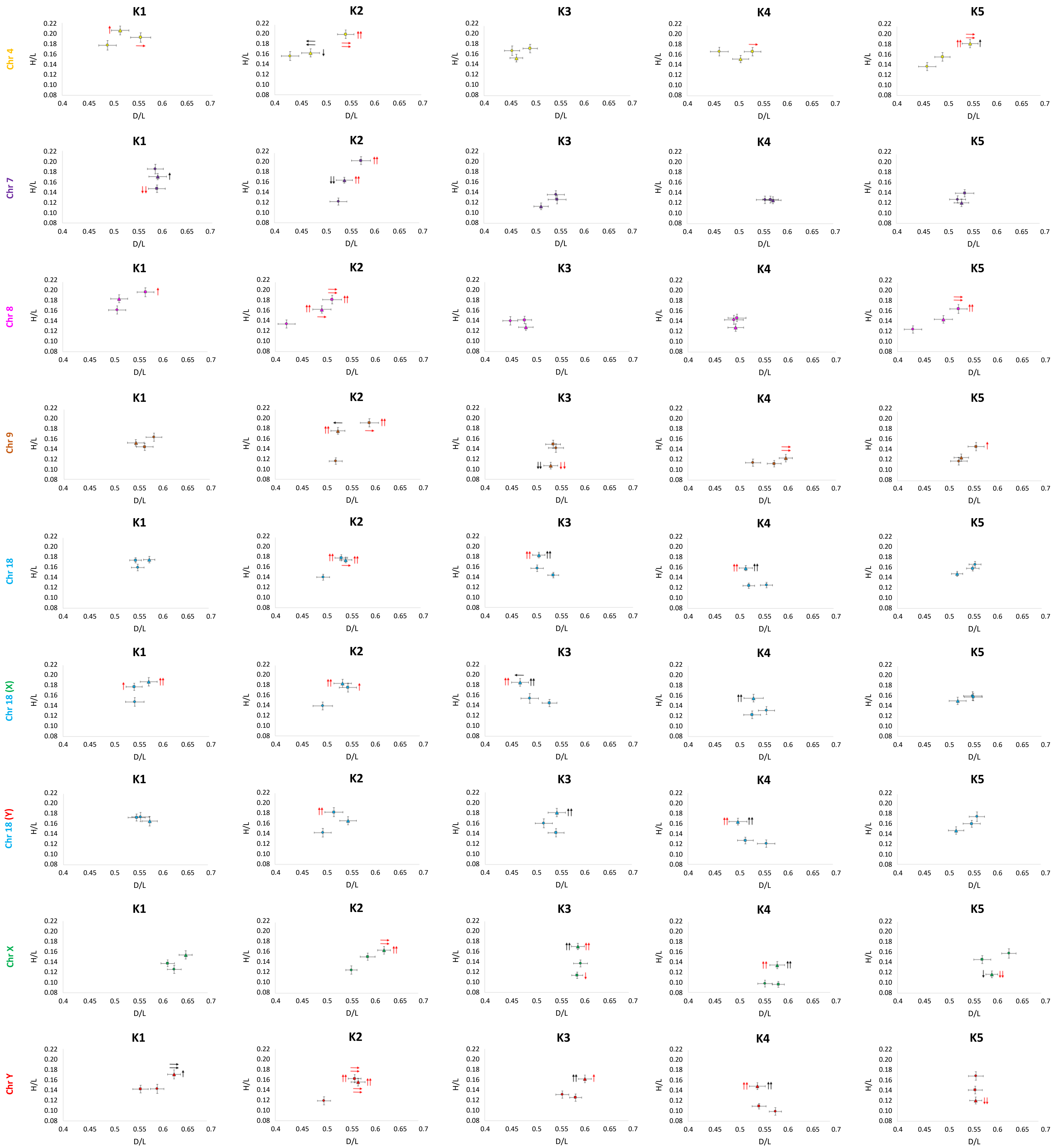
