## Supplementary material for "Impact of Sperm Fractionation on Chromosome Positioning, Chromatin Integrity, DNA Methylation and Hydroxymethylation Level": List of Additional files

Additional file 1 .pdf FISH probes used for experiments (Cytocell, Cambridge, UK).

Additional file 2 .xlsx Individual data for results of sperm chromatin integrity evaluation in non-fractionated sperm population (native ejaculate) and good-quality fractions (swim up fraction (SU) and bottom layer of Percoll gradient (BPG)) in each evaluated case.

Additional file 3 .xlsx Individual data for linear positioning of centromeres of chromosomes: 4, 7, 8, 9, 18, X and Y in spermatozoa in each evaluated case.

Additional file 4 .xlsx Individual data for radial positioning of centromeres of chromosomes: 4, 7, 8, 9, 18, X and Y in spermatozoa in each evaluated case.

Additional file 5 .pdf Radial positioning of the each of examined centromeres (4, 7, 8, 9, 18, X, Y) separately within the sperm nucleus in non-fractionated ejaculate, swim up fraction (SU) and Percoll gradient (BPG), according to data in Additional file 3 (circle: non fractionated ejaculate, triangle: swim up fraction, square: bottom layer of Percoll gradient). Localizations that differ significantly from the non-fractionated ejaculate mean value were indicated by red arrows: double arrow for p ≤ 0.01, single arrow for p ≤ 0.05. Statistically significant differences between SU and BPG fractions were indicated by black arrows: double arrow for p ≤ 0.01, single arrow for 0.01 < p ≤ 0.05. Arrows also indicate the direction of the observed shift (repositioning) of centromeres. Bars show standard errors (SE).

Additional file 6 .pdf Individual data for radial positioning of the examined centromeres (4, 7, 8, 9, 18, X, Y) within the sperm nucleus in non-fractionated ejaculate, swim up fraction (SU) and Percoll gradient (BPG) in each evaluated case, according to the data in Additional file 3 (circle: non fractionated ejaculate, triangle: swim up fraction, square: bottom layer of Percoll gradient). The mean control values are marked in black. Red asterisks indicate statistically significant differences compared to the mean values. Bars show standard errors (SE).

Additional file 7 .xlsx Distances between the centromeres of chromosomal pairs: 4 and 8, 7 and 9, 18 and X, 18 and Y in non-fractionated ejaculate and good-quality fractions (swim up fraction (SU) and bottom layer of Percoll gradient (BPG)) in each evaluated case.
